# The molecular asynchrony of single cells

**DOI:** 10.64898/2026.06.02.729594

**Authors:** Boshi Fu, Robert Tan, Zhenkun Cao, Xiuzhen Bai, Dongsheng Bai, Jinghui Song, Chenxu Zhu

## Abstract

Integrating single-cell transcriptomic and epigenomic data provides a robust framework for investigating gene regulation mechanisms. Existing analyses typically treat these modalities as synchronized features that can be translated in a static manner; however, the temporal delays that underpin cellular kinetics are intrinsic to dynamic biological systems. To address this limitation, we propose utilizing the “molecular asynchrony” within regulatory hierarchies to determine the thermodynamic properties of individual cells. Here, we present SeqTag, a single-cell multiomics sequencing method that simultaneously profiles the transcriptome, chromatin accessibility, and histone modifications, supported by an analytical framework to identify asynchronous states across regulatory layers for characterization of single-cell kinetics. By measuring the epigenetic priming potential and remodeling rates during adult mouse oligodendrogenesis, we delineated a sequential program for bivalency resolution as maturing cells traverse Waddington’s landscape. This process becomes increasingly decoupled with age, a change linked to a drift in progenitor cell-fate probabilities. By identifying entropy-driving regulatory elements, we characterized the aging-related decline in cell identity across various cell types and proposed a dynamic model linking static genetic variants to the risk of late-onset diseases. In summary, our integrated approach established a unified framework for employing multimodal single-cell genomics to model the kinetics of complex cellular processes.

## Introduction

The Waddington’s epigenetic landscape conceptualizes cells as marbles traveling through a topographically complex surface, where valleys denote stable cell states, and ridges represent developmental barriers^1^. It is widely recognized that cell identity is highly dynamic, responding to various developmental stimuli as cells traverse the landscape^2^. In fact, cells not only vary in their phenotypic identity but also differ in their energetic states. At the macroscopic level, a functionally relevant cellular population (e.g., a biological tissue) is a complex ensemble of cells partitioned over various thermodynamic states. For example, self-renewing progenitors exist in high-entropy steady states, whereas committed progenitors are in a metastable state that gathers epigenetic priming energy. Once the energy threshold is surpassed, cells transition to high-energy intermediates, accompanied by rapid changes in phenotypic states (e.g., differentiation). Even in terminal states, where mature cells are at equilibrium, they may undergo a gradual entropic decline, leading to processes such as the loss of cell identity during aging^3^. Therefore, characterizing the thermodynamic parameters of individual cells is necessary for identifying both the factors that support the differentiation and homeostasis of healthy tissues and the energetic barriers that fail during disease and aging.

Advancements in single-cell and spatial genomics have enabled the digital characterization of complex cellular populations; however, most measurements capture static snapshots of cells^4^. This presents the challenge of discerning whether a cell is in a resting, steady state or is actively undergoing a state transition. To address this challenge, innovations were implemented, advancing single-cell analysis from descriptive to quantitative and predictive frameworks. For example, by ordering cells according to a mathematically derived “pseudotime” coordinate based on transcriptomic similarity, methodologies have been devised to elucidate the continuum and branching dynamics of cell-state transitions^5-7^. Nonetheless, distance-based approaches lack a thermodynamic foundation, which characterizes the pathway without addressing the underlying driving forces. RNA velocity incorporates biophysical properties of mRNA, utilizing the ratio of spliced and unspliced mRNA to predict the immediate future of individual cells^8-10^. However, the dependency on fast kinetic processes limits its applicability to slow-timescale processes (e.g., drug treatment outcomes). Optimal transport calculates the most probabilistically efficient paths cells follow over time^11^; however, it requires dense, sequential data sampling, and applying it to inherently asynchronous processes (e.g., mammalian aging) is challenging. Causal regulatory frameworks, including SCENIC and Chromatin Potential, leverage multimodal single-cell datasets to provide mechanistic insights^12,13^. We previously explored leveraging cytosine modification entropy, which quantifies the complexity of relationships between DNA methylation and active demethylation, to detect cells undergoing active DNA methylation reprogramming^14^. Although they are effective for modeling cells undergoing active phenotypic state changes, characterizing mixed cellular populations spanning a continuum of thermodynamic states remains challenging.

In chemical thermodynamics, a reaction is described as a substrate being transformed into products, passing through high-energy intermediate states along the reaction coordinate^15^. In the context of single-cell kinetics, the ergodic hypothesis^16^ can be reinterpreted as the idea that the observed asynchronous cells at a single moment are mathematically equivalent to observing a single cell’s probability distribution over a long time. According to the Boltzmann distribution, the density of cells observed in a specific cellular state (probability) is inversely, exponentially proportional to the “energy” of the state^17^. Therefore, the underlying “potential energy surface” of Waddington’s landscape can be inferred from highly sampled single-cell data. However, cellular state transitions are not instantaneous across all molecular layers, and inferring such temporal lag from independently generated datasets is challenging. For example, a significant limitation in single-cell genomics is technical sparsity, which considerably constrains the accuracy of 1-to-1 cross-modal mapping. Therefore, multimodal single-cell technologies that experimentally link distinct molecular layers are essential to identifying cellular priming. Nonetheless, being epigenetically primed does not guarantee that a given cell will spontaneously undergo a state transition (e.g., in immune cells, early-response genes are epigenetically primed but activate only upon receiving external signals)^18^. In fact, such asynchrony exists not only between the epigenome and transcriptome but also between different epigenetic layers. For example, changes in chromatin accessibility usually precede or coincide with changes in the polycomb-associated repressive histone mark H3K27me3^19,20^. This disequilibrium is not incidental noise; rather, it serves as a direct proxy for the kinetic properties of individual cells, which can be leveraged to model cellular thermodynamic parameters and enable quantitative predictions of future states.

To capture this transient disequilibrium, it is necessary to measure the distinct regulatory layers simultaneously. Chromatin accessibility reflects an ensemble state of epigenetic permissiveness, whereas specific histone marks give detailed biochemical instructions. Joint profiling of these two layers, along with the transcriptome from the same cell, is essential for linking the “leading” and “lagging” epigenetic states to a realized phenotypic state. Therefore, we developed SeqTag, a scalable technique for tri-modal analysis of ATAC-seq, CUT&Tag, and RNA-seq in single cells. We analyzed the mouse cerebral cortex during aging, a native system that contains various energetic states, including self-renewing and committed progenitors, proliferating glial cells, and mature neurons from different ages. Our results reveal that aging manifests as a flattening of the epigenetic priming energy barrier, thereby diminishing regulatory fidelity in both mature neurons and dynamic populations such as oligodendrocyte precursor cells (OPCs). By utilizing chromatin accessibility as an “internal clock”, we estimated the reprogramming rates of histone modifications H3K27ac and H3K27me3 and found that the two processes become decoupled and correlate with a drift in cell-fate probability in aged cells. Moreover, quantitative parameters help assess how individual genes and regulatory elements contribute to changes in cellular behavior, thereby aiding the identification of potential intervention targets. Thus, modeling cellular kinetics via multimodal molecular asynchrony uncovers regulatory program changes that are hidden by current static analysis approaches.

## Results

### The thermodynamics model of single cells

To understand the cellular kinetics embedded in the trimodal dataset (chromatin accessibility, histone modifications, and the transcriptome), we conceptualized state transitions through non-equilibrium thermodynamics and transition-state theory^21-23^. In this regime, the epigenome dictates the “potential energy surface”, and the transcriptome acts as the lagging functional readout. By co-projecting multimodal single-cell data (e.g., chromatin accessibility, histone marks, transcriptome) into a unified latent space, the “temporal lag” between molecular layers can be measured (**Figure 1A**). These asynchronies can be exploited to define three thermodynamic parameters for individual cells. Priming Energy is defined as the preloaded epigenetic changes, reflected in the cell’s chromatin accessibility (the “leading” state), which are prepared for the subsequent histone modification (the “lagging” state) to catch up in the near future. This metric isolates the energy available from the synchronized actions of distinct epigenetic layers to overcome reprogramming barriers, based on the cellular distributions in snapshot single-cell data. To estimate how quickly cells move between successive states, we define the Epigenetic Remodeling Rate as the distance between the “leading” and “lagging” epigenetic states that the cell must traverse in a short timeframe. Finally, to quantify a cell’s intrinsic epigenetic plasticity, we calculated Regulatory Entropy across different epigenetic modalities, based on the range of possible phenotypic states within its immediate epigenetic environment (**Methods**).

**Figure 1.**
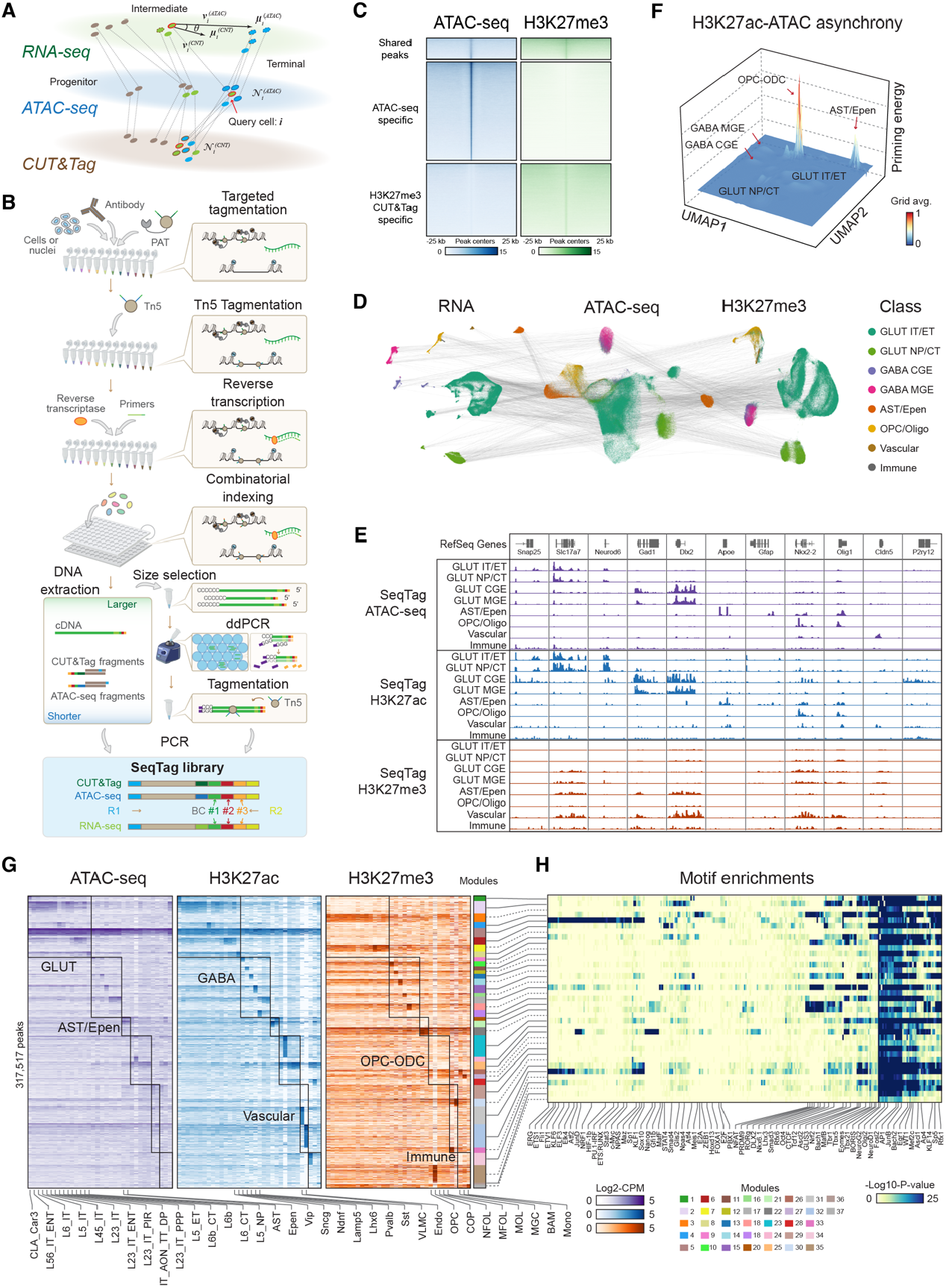
SeqTag unveils the thermodynamics landscape of the brain. **(A)** The displacement vectors of cells can be acquired through cross-modality nearest neighbors, thereby facilitating the quantification of multimodal molecular asynchrony. **(B)** Schematics of SeqTag. Antibody-targeted and non-targeted tagementation steps were sequentially performed to minimize signal leakage. **(C)** Heatmap showing the detected signals within shared, ATAC-seq specific, and H3K27me3 specific peak regions derived from SeqTag profiles. **(D)** UMAP showing the embedding of single cells derived from gene expression, chromatin accessibility, and H3K27me3 data of SeqTag mouse cerebral cortex samples. Each dot represents an individual cell and is color-coded according to its L1 class. The same cells across the three embeddings are connected by grey lines. **(E)** SeqTag DNA signals of representative marker gene loci for major brain cell classes. **(F)** The 3-D UMAP showing the priming energy calculated from H3K27ac-ATAC-seq asynchrony. The x and y axes represent UMAP embedding coordinates derived from gene expression data, while the z-axis depicts the grid-averaged priming energy values. **(G)** Heatmaps displaying chromatin accessibility, along with histone H3K27ac and H3K27me3 signals on the identified cCREs across various subclasses. Module memberships of cCREs are shown beside. **(H)** Heatmap showing the motif enrichment scores of different cCRE modules.

### The SeqTag technology

We designed a sequential tagmentation approach (SeqTag) to simultaneously measure chromatin accessibility^24^ and histone modifications^24^, alongside the transcriptome^25^, in the same single cells (**Figure 1B, Methods**). In the SeqTag procedure, cells or nuclei are permeabilized, followed by antibody-targeted tagmentation by protein A-Tn5 (PAT) fusion protein^26^ to attach the first set of barcoded oligo DNA adaptors. Next, cells or nuclei are washed to remove residual PAT, followed by the 2nd tagmentation step with Tn5 to attach oligo DNA adaptors to accessible chromatin regions. Reverse transcription (RT) is then performed with the barcoded random hexamer and oligo-dT primers. These three sets of oligo DNA barcodes share the same tube-specific sequence and an identity handle sequence, enabling simultaneous combinatorial indexing^25^. Finally, the barcoded cells or nuclei are lysed to purify cDNA and chromatin fragments, and DNA and RNA libraries are independently amplified and sequenced. The PAT and Tn5 adaptors contain a unique modality-specific sequence that enables computational signal demultiplexing.

Since capturing two chromatin modalities relies on a similar Tn5-based tagmentation, which has been shown to risk barcode exchange^27^. We performed the two transposition reactions in temporally separated compartments^27^, thereby minimizing cross-modality signal leakage (**Figure 1C**). We first validated the specificity of chromatin detection on HeLa cells: profiles obtained from sequential tagmentation showed good agreement with independently generated single-modality reference data, and displayed a high signal-to-noise ratio (**Figures S1A, B**). Microfluidics-based scRNA-seq assays frequently showed higher sensitivity than plate-based combinatorial barcoding^28^, possibly due to pre-amplification steps within microcompartments that reduced amplification bias, thereby increasing the recovery of low-abundance transcripts. We designed an instrument-free microdroplet-based^29^ pre-amplification step to distribute a limited number of cDNA molecules into each microcompartment and increased cDNA library complexity by > 20% (**Figure S1C**). Next, we performed species-mixing experiments (human HeLa and mouse NIH/3T3 cells) to validate SeqTag in single-cell analysis. Among the barcodes with sufficient numbers of reads in all three modalities, 2,827 out of 3,011 barcodes can be uniquely assigned to a single species, which is comparable with previous combinatorial indexing single-cell assays^25,30,31^ (**Figure S1D**). We detected median numbers per cell of >12,000 transcripts and 976 to >6,000 unique chromatin fragments for the ATAC-seq, H3K27ac, and H3K27me3 modalities in HeLa cells (**Figure S1E**). These data support that SeqTag simultaneously measures chromatin accessibility, histone modifications, and the transcriptome in single cells with high sensitivity and specificity.

### Single-cell tri-modal atlas of the mouse cerebral cortex

To demonstrate the utility of SeqTag for analyzing heterogeneous cellular populations, we generated single-cell profiles from the cerebral cortex of mice at three ages. Antibodies targeting two major histone marks, the active mark H3K27ac and the repressive mark H3K27me3, were used for the CUT&Tag modality. SeqTag captures median numbers of 14,930 and 19,027 transcripts and 14,554 and 19,316 chromatin fragments per cell for H3K27ac and H3K27me3 SeqTag experiments, respectively, which are comparable to or higher than those from existing dual-omics methods^13,32-34^ (**Figure S1F-H**). By sequencing additional libraries to moderate depth (average 50,000 reads per cell) and filtering low-coverage barcodes, we obtained multiomics profiles for 450,031 cells (217,501 with H3K27ac profiles and 232,530 with H3K27me3 profiles).

Independent cell clustering was performed based on transcriptome, chromatin accessibility, and histone modification modalities, and the results agreed well with each other and the reference Allen Institute mouse brain atlas^35^ (**Figures 1D-F** and **S1I**). We grouped and annotated the cells into 8 primary classes (L1 classes) and 34 cell subclasses (L2 subclasses) according to the reference atlas of the relevant brain region^35^ (**Figures 1E** and **S1J**). Next, we performed peak calling with MACS3^36^ on chromatin accessibility profiles and identified 317,517 candidate cis-regulatory elements (cCREs), which were further grouped into 37 modules according to the signal combinations from aggregated chromatin accessibility, active and repressive histone marks across the 34 subclasses using non-negative matrix factorization (**Figure 1G** and **Table S4**). Motif enrichment analysis of the cCRE modules robustly identified the key regulators in each brain cell lineage, including SOX10 for oligodendrogenesis^37^ (module 23), DLX2 for GABAergic neurons^38^ (module 17), and Stat3 for microglia cells^39^ (module 34) (**Figure 1H**). Thus, SeqTag provided a comprehensive reference map of the mouse brain epigenome and transcriptome.

**Figure S1.**
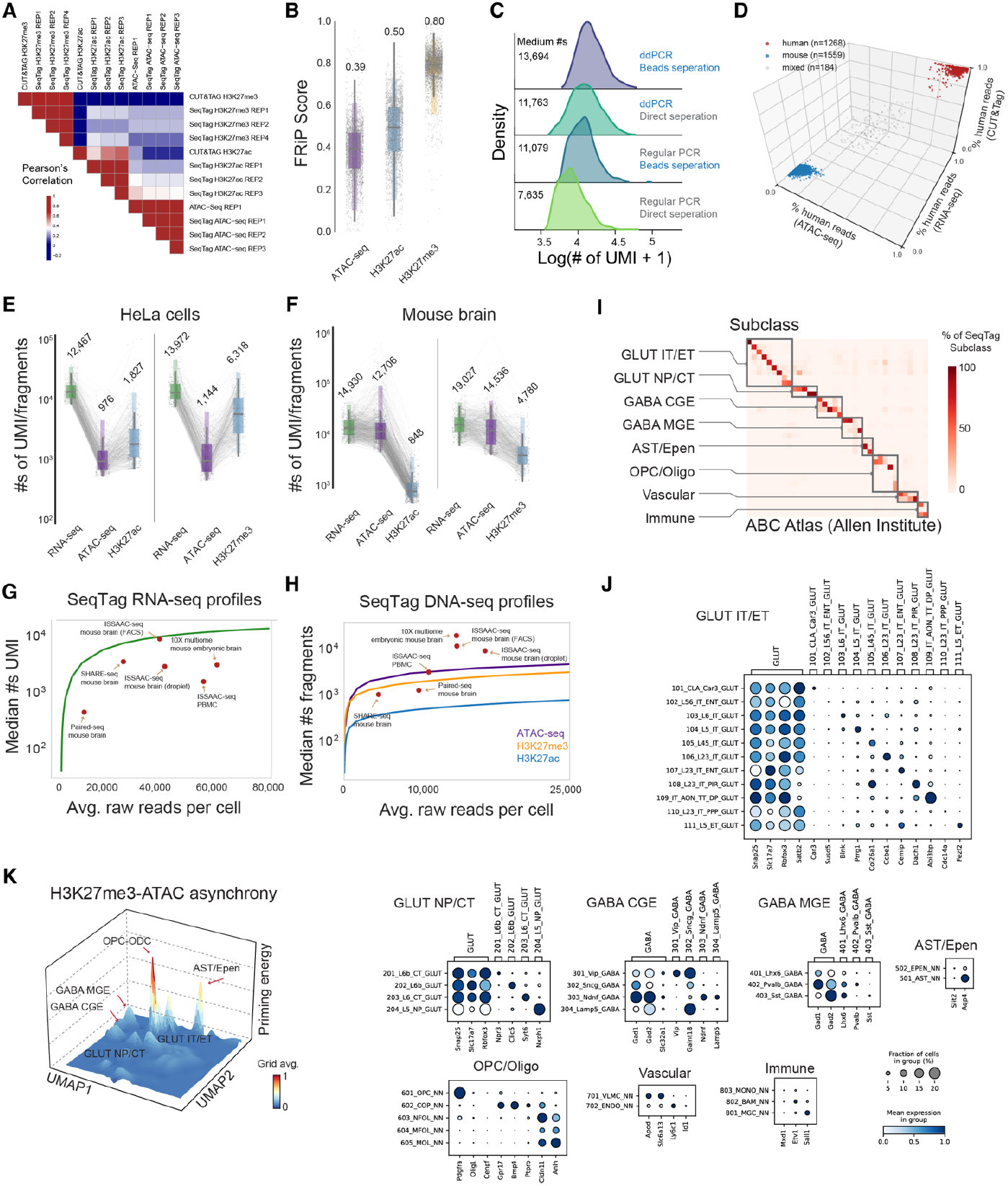
Validation of SeqTag technology. **(A)** Heatmap showing the Pearson correlation coefficients among H3K27ac CUT&Tag, H3K27me3 CUT&Tag, ATAC-seq, and SeqTag H3K27ac, H3K27me3, and ATAC-seq profiles derived from HeLa cells. **(B)** Boxplot showing the FRiP scores for SeqTag ATAC-seq, H3K27ac, and H3K27me3 profiles generated from HeLa cells. The median scores are annotated above the plot. **(C)** Ridge plots showing the number of transcripts captured per cell from different library amplification methods. All conditions were downsampled to ensure an identical average sequencing depth per cell. **(D)** 3-D scatter plot showing the barcode collision resulting from species-mixing experiments. **(E), (F)** Boxplot showing the library complexities of deep-sequenced SeqTag libraries derived from HeLa cells (E) and mouse cerebral cortex (F). **(G), (H)** The saturation plot showing the library complexities of SeqTag RNA (G) and DNA (H) profiles at various sequencing depths. Public datasets are indicated for comparative purposes. **(I)** Heatmap showing the confusion matrix for RNA-based independent clustering and reference mapping utilizing the ABC Atlas. **(J)** Dot plots showing the gene expression levels of L2 subclasses across the eight major classes. **(K)** 3-D UMAP showing the priming energy computed from H3K27me3-ATAC-seq asynchrony. The x and y axes represent UMAP embedding coordinates derived from gene expression data, while the z axis indicates the grid-averaged priming energy values.

### The epigenetic priming energy landscape of mouse brain cells

The adult brain contains complex cell populations that not only differ in their identities but also occupy distinct transition states. For each cell, we identified its k-nearest neighbors across the three molecular layers and projected them from the two epigenomic manifolds back into the RNA embeddings, yielding two priming vectors (RNA-to-ATAC and RNA-to-CUT&Tag). The discrepancies between the transcriptome and epigenome reflect the gradients of local thermodynamic forces that pull the cell toward new states, and these gradients are proportional to the magnitude and coherence of priming vectors. The magnitude corresponds to the difference in norms between the RNA-to-ATAC and RNA-to-CUT&Tag priming vectors, representing the tension of “hidden” epigenetic layers as they transition from the “observed” histone modification state to the chromatin accessibility ensemble profile. Coherence is assessed by the cosine similarity between the two priming vectors, ensuring the forces guiding the cell’s state change are directionally aligned. A pseudo-RNA-to-RNA priming vector is used to account for basal stochasticity and technical variation (**Figure 1A**). As expected, based on the H3K27ac-ATAC asynchrony, cells within the OPC-to-ODC trajectory displayed the highest priming energy, followed by astrocytes that retain the ability for reactive astrogliosis^40^; on the other hand, all neuron cell types exhibit a basal level of priming energy (**Figure 1F**). We also observed a similar trend from the H3K27me3-ATAC asynchrony (**Figure S1K**). Thus, the epigenetic priming energy of both active and repressive histone marks pinpoints cells actively undergoing state transitions.

### Bivalency resolution during oligodendrogenesis

To investigate how epigenetic priming energy changes during cell-state transitions, we isolated oligodendrocyte lineage cells and performed pseudotime analysis (**Figures 2A, S2A**, and **S2B**). The cells’ diffusion pseudotime scores are highly correlated with their PC1 scores, reflecting the molecular continuum from proliferating OPCs to mature ODCs (**Figure S2A**). Since changes in chromatin accessibility usually precede or are concordant with the changes in histone modifications, the gap between the epigenetic layers can inform the direction of state transition without the requirement of prior knowledge (**Figure 2B**). We then calculated the average priming energy scores for cells with similar pseudotime and reconstructed the energy landscape along the differentiation coordinate (**Figures 2C, S2C**, and **S2D**). The two histone marks showed different but overall similar trends: cells rapidly lose priming energy as they transition from OPCs to immature ODCs. Thus, the remodeling of active and repressive chromatin during ODC maturation follows a coupled but asynchronous program.

**Figure 2.**
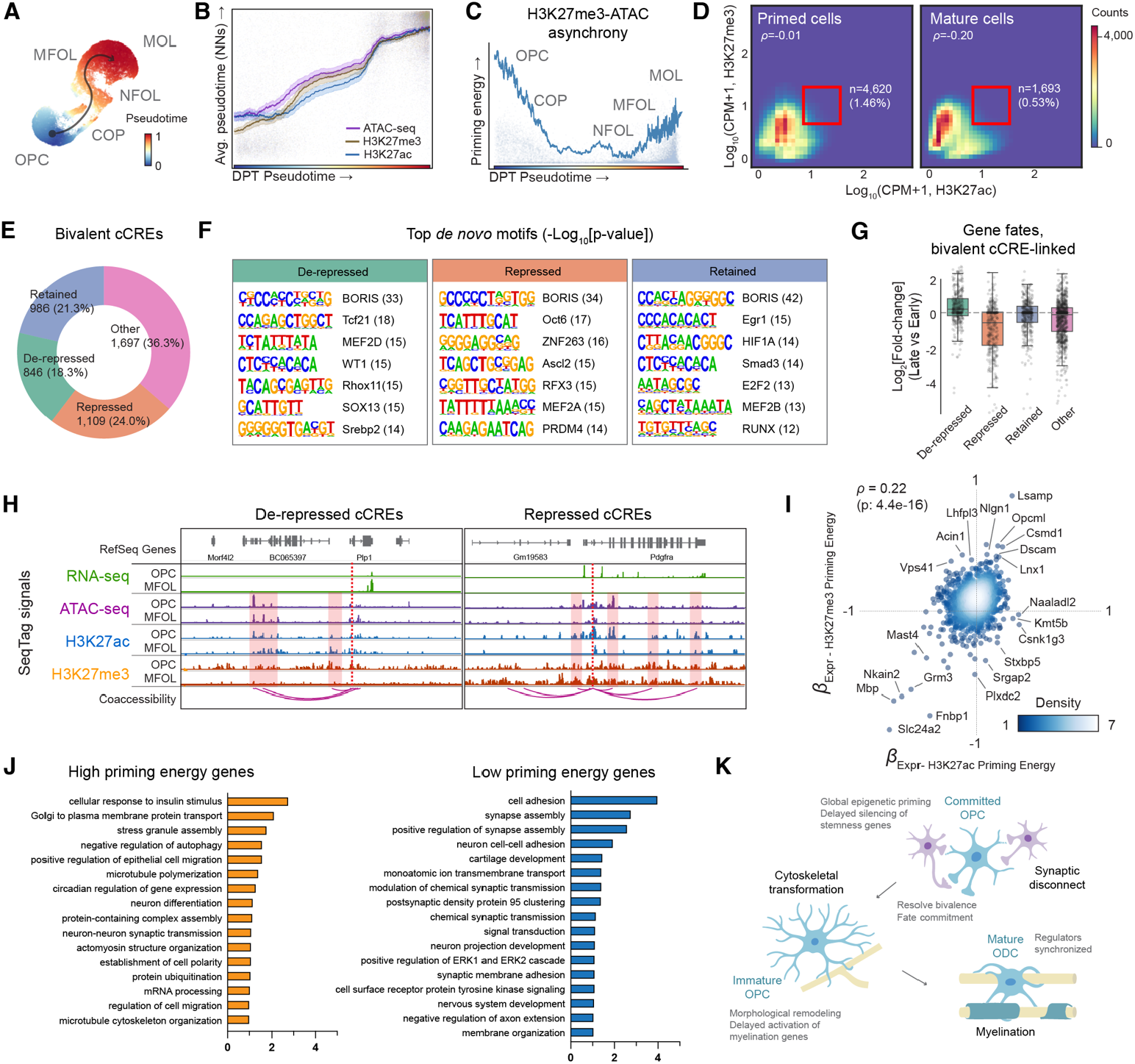
Asynchronous chromatin remodeling during oligodendrocyte maturation. **(A)** UMAP showing the trajectory of OPC differentiation. Cells are color-coded based on diffusion pseudotime scores. **(B)** Line plots showing the temporal relationships of chromatin accessibility, along with active and repressive histone marks reprogramming during OPC differentiation. **(C)** Priming energy landscape derived from H3K27me3-ATAC asynchrony. **(D)** Heatmap showing the relationships between H3K27me3 and H3K27ac modification levels in primed and mature cells. **(E)** Fraction of bivalent cCREs in primed cells retained, lose H3K27me3, or lost H3K27ac in mature cells. Bivalent cCREs that lose both H3K27me3 and H3K27ac were grouped as Other. **(F)** Top enriched de novo motifs for de-repressed, repressed, and retained bivalent cCREs. **(G)** Boxplots showing the variations in transcription levels during OPC differentiation for genes linked to different groups of bivalent cCREs. **(H)** Genome browser view showing the representative loci of de-repressed and repressed bivalent cCREs. **(I)** Scatter plot showing the relationships between gene expression levels and the priming energy of H3K27ac and H3K27me3. **(J)** Top enriched GO terms for high priming energy and low priming energy genes. **(K)** Proposed model of the sequential epigenetic remodeling during OPC maturation.

**Figure S2.**
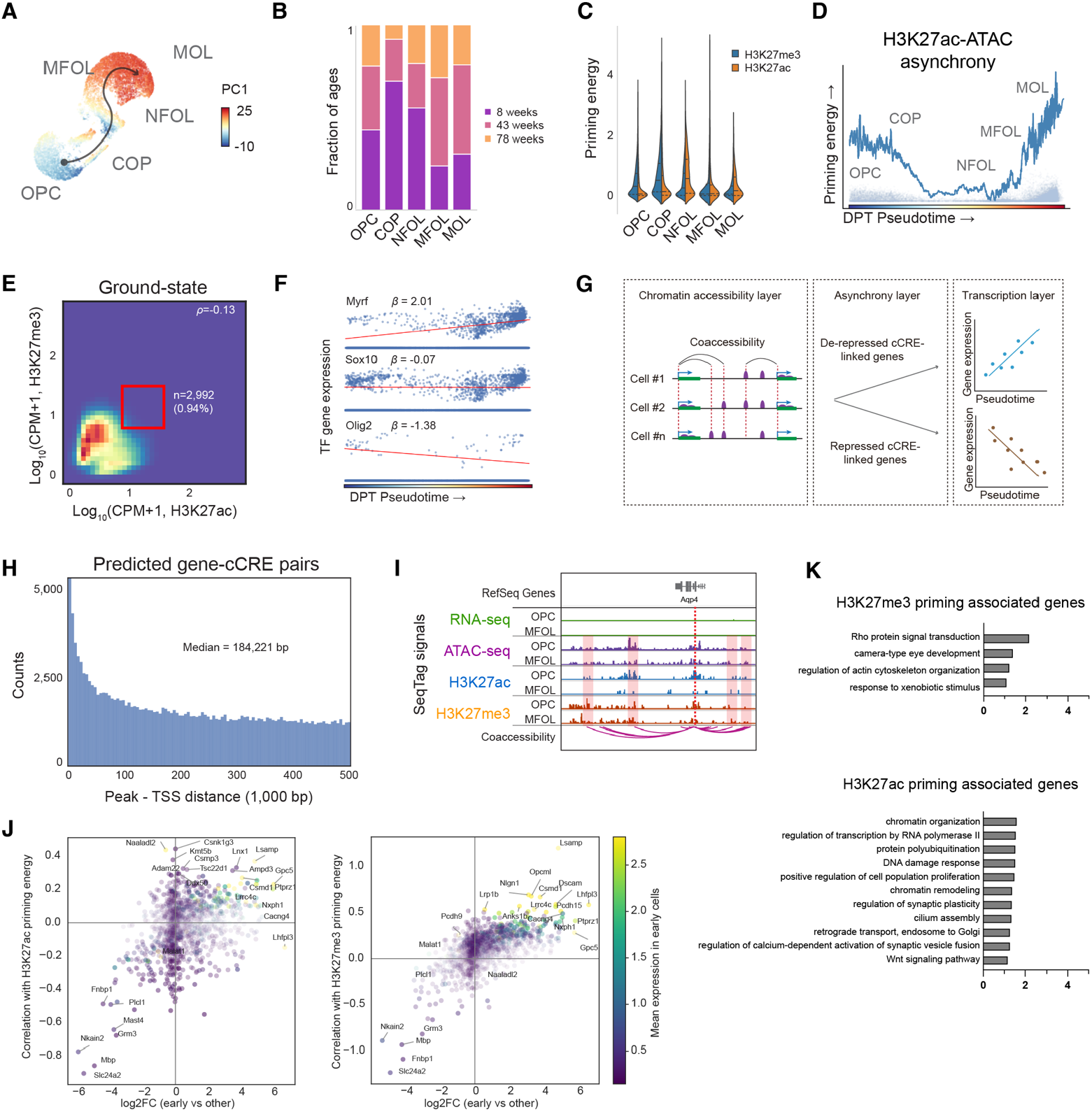
Asynchronous epigenetic remodeling of oligodendrocyte maturation. **(A)** UMAP showing the trajectory of OPC differentiation. Cells are indicated by colors corresponding to PC1 scores. **(B)** Stacked barplots showing the proportion of cells from various age groups for OPC, immature, and mature ODCs. **(C)** Violin plots showing each cell’s epigenetic priming energy calculated from H3K27ac-ATAC and H3K27me3-ATAC asynchrony. **(D)** Priming energy landscape derived from H3K27ac-ATAC asynchrony. **(E)** Heatmap showing the relationships between H3K27me3 and H3K27ac modification levels in ground state immature cells. **(F)** Representative TF gene expression changes during OPC differentiation. **(G)** Schematics of cCRE-gene pairs prediction from the multilayered regulatory profiles. **(H)** Histogram showing the distributions of distances between cCREs and TSSs of predicted target genes. **(I)** Genome browser view of the Aqp4 locus and the repressed bivalent cCREs. **(J)** Scatter plots showing the relationships between gene-priming energy associations and gene expression level changes during ODC maturation. Each data point signifies a gene and is color-coded based on its expression level in immature cells. Left: H3K27ac priming energy, right: H3K27me3 priming energy. **(K)** Top enriched GO terms for genes associated with H3K27me3 and H3K27ac priming energies.

We observed that the priming energy of OPCs follows a bimodal distribution (**Figure S2C**). The “high-energy” group likely consists of primed OPCs that are actively preparing to overcome the activation energy barrier for differentiation and myelination; the “low-energy” majority, by contrast, remains in a ground state to preserve the progenitor reservoir. To test this, we classified the immature cells (DPT pseudotime < 0.75) into the primed state (priming energy > 0.2) and the ground-state (priming energy < 0.1), and compared their genome-wide H3K27ac and H3K27me3 modification levels (**Figures 2D, S2E**). As expected, compared to mature cells (DPT pseudotime > 0.75) and ground-state cells, the primed cells exhibit less negative correlation in the modification levels of the two histone marks with opposite regulatory functions, and a higher fraction of the bivalent cCREs as defined by the prevalence of both marks (Odds ratio: 2.75 vs mature, and 1.55 vs ground-state progenitor cells). As the cells mature, over one-third of the bivalent cCREs lost both H3K27 acetylation and methylation. Among the remaining, 24.0% lost the activation mark and became fully repressed, 18.3% demethylated to activate, and the rest remained bivalently marked (**Figure 2E**). The binding motifs of WT1 and the SOX family protein were found to be significantly enriched in de-repressed cCREs (**Figure 2F**). WT1 is a master transcription factor that binds to GC-rich regions of poised cCREs and modulates the resolution of bivalent states^41^, together with Sox10, another key regulator for oligodendrogenesis^37^, to trigger epigenetic priming and subsequent myelination (**Figure S2F**). To explore the regulatory consequences of this bivalency resolution, we predicted the target genes of the detected cCREs according to gene-cCRE co-accessibility in the ATAC-seq modality (**Figures S2G-I**). For linked genes across different cCRE groups, the de-repressed bivalent cCRE targets showed an evaluated overall expression level in mature ODCs, whereas the other groups remained unchanged or were silenced (**Figures 2G, 2H**). Together, our data suggest that molecular asynchrony reveals a bidirectional process of chromatin remodeling that leads to functional outcomes.

### Sequential epigenetic remodeling of oligodendrogenesis

Trajectory inference and pseudotime analysis are powerful tools to provide regulatory insights by uncovering the hidden continuum in single-cell data^5-7^. However, such analysis relies on an idealized, post hoc consensus trajectory; regressing gene expression against it failed to capture the rate-limiting intermediate asynchronous states and is incapable of distinguishing kinetic drivers from passenger genes^17^. Our molecular asynchrony model estimates epigenetic priming potential based on the non-equilibrium geometric differences across regulatory layers. We suggest that this analysis could provide the thermodynamic resolution needed to pinpoint key regulatory factors that can overcome intermediate barrier states. To test this, we performed linear logistic regression of gene expression levels on single-cell priming energies and identified both well-known and less-studied regulators (e.g., Mbp, Kmt5b, Lsamp). Note that not all top-ranked genes are identifiable through differential analysis across maturation stages, as these genes may serve as both drivers and outcomes of the differentiation process (**Figure S2J**). According to their associations with remodeling of H3K27ac and H3K27me3, we classified the top-ranked genes into four groups: (1) group A: bivalent remodeling genes; (2) group B: H3K27ac-priming associated genes; (3) H3K27me3-priming associated genes; and (4) group D: genes negatively associated with priming (**Figures 2I, 2J, S2K**).

Our data showed that immediately after chromatin accessibility was reprogrammed, H3K27me3 modification began to catch up with the leading state, while H3K27ac remodeling experienced a brief delay before all three modalities became fully synchronized in mature cells (**Figure 2B**). Based on this, we proposed a sequential model of epigenetic remodeling during oligodendrogenesis (**Figure 2K**). This process starts with pioneer factors (e.g., Sox10, Olig1/2) that decondense closed chromatin and establish the open chromatin blueprint for subsequent histone modifications remodeling. The following step is H3K27me3 priming, which triggers cell recycling pathways (e.g., group C genes like Vps41, essential for late-endosome-to-lysosome fusion and autophagy^42^, and Acin1, involved in RNA processing and chromatin condensation^43^). This prepares cells for metabolic and structural challenges as they exit the stable stemness state. The next step involves resolving bivalency, which either activates or represses group A genes involved in synapse organization and cell adhesion (e.g., Lsamp, Opcml, Nlgn1, and Dscam). This agrees with the fact that OPCs receive synaptic inputs from neurons, which must be dismantled before they mature into myelinating oligodendrocytes^44^. Meanwhile, group B genes are activated to facilitate morphological remodeling (e.g., Srgap2 regulates actin dynamics^45^, and Stxbp5 governs SNARE-mediated vesicle fusion and exocytosis^46^) and to prepare for the downregulation of stemness genes, thereby synchronizing regulatory layers as the cell matures (e.g., Kmt5b, a histone H4K20 methyltransferase). This stepwise process for resolving regulatory conflicts helps the cell to overcome the activation energy barrier to reach a fully synchronized, stable, mature state.

### Aging decouples the remodeling of different histone modifications

We next asked how the thermodynamic properties of OPC epigenetic remodeling are influenced by the process of biological aging. In aged cells, both H3K27ac and H3K27me3 exhibited reduced baseline-level priming energy in NFOL and MFOL, indicating a higher energy barrier to achieve complete myelination (**Figure S3A**). Interestingly, an energy trap was observed in early-committed OPCs of aged mice, appearing as a metastable intermediate state midway through maturation. As cells traverse the epigenetic landscape, this trap does not alter the overall direction of chromatin remodeling; however, the effects on remodeling rates across different epigenetic layers may differ, potentially decoupling overall regulatory conflict resolution. To test this, it is necessary to directly compare the remodeling kinetics of various histone marks. Because chromatin accessibility reflects the combined effects of multiple chromatin modifications that influence chromatin structure, SeqTag data is well-suited for this purpose by projecting different histone marks from separate latent spaces onto a common coordinate system, leveraging the ATAC-seq profile as an internal reference.

To estimate the rate of histone mark remodeling for individual cells, we determine its transition distribution by finding its k-nearest neighbors in the lagging modality (CUT&Tag) and projecting these neighbors onto the leading modality (ATAC-seq) embeddings (**Figure 3A**). The topological segmentation of these projected neighbors captures the temporal imbalance in high-probability microstates during an active transition, in which some cells remain anchored to the current state, and others leap toward the future state. We fit a bimodal distribution to the locally scaled neighbor distances, and the distance between the distribution’s two density maxima indicates how much the lagging modality needs to catch up with the leading modality within the next time interval (**Figures 3B, 3C**, and **S3B**). We found that the differences in average H3K27ac and H3K27me3 remodeling rates vary in cell maturation, with early-committed cells exhibiting considerably faster H3K27me3 reprogramming (**Figure S3C**). However, these differences likely result from the abnormal surge in H3K27me3 changes in aged cells during the early stages of differentiation (**Figure 3D**). We performed linear regression analysis of cCRE H3K27me3 levels against their remodeling rates. The results showed that the top associated elements are frequently hypermethylated in committed oligodendrocyte precursors (COPs) but tend to lose this signature with age (**Figure 3E-G**). On the other hand, H3K27ac remodeling did not show this trend (**Figures S3D-F**). Interestingly, the H3K27me3 remodeling-associated cCREs are enriched not only in the COP hypermethylated region (M36), but also in astrocyte-specific H3K27ac-modified regions (M21) (**Figures 3H** and **S3G**). For example, the lineage switch Prdm6 directs stem cells toward neuronal and astroglial pathways^47^, and in OPCs, its transcription is repressed by H3K27me3; however, it loses this repressive mark with age (**Figure 3I**). While this methylation erosion in aged OPCs does not trigger Prdm16 activation, de-repressing this gene could increase the risk of driving older cells into a pro-stemness state^47^, potentially contributing to glioma initiation^48^. Thus, molecular asynchrony provides the kinetic resolution to reveal temporal relationships in the remodeling of various epigenetic layers.

**Figure 3.**
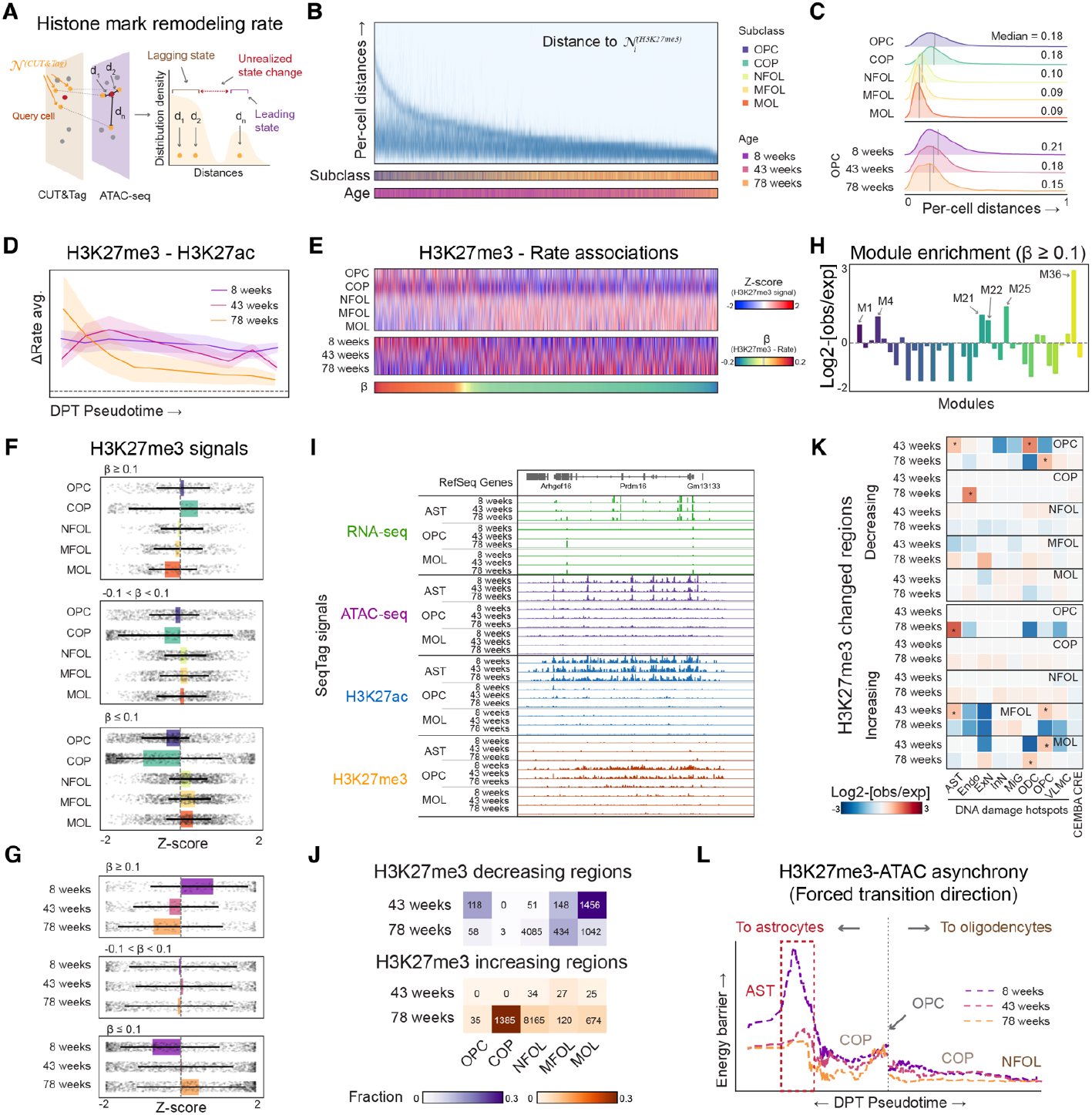
Decoupled epigenetic remodeling in aged OPCs linked to fate probability drift. **(A)** The rate of histone mark remodeling in cells can be estimated by identifying the bimodal distribution of their nearest neighbors within the chromatin accessibility embedding space. **(B)** Heatmap showing the normalized per-cell distance between lagging and leading states of individual cells derived from H3K27me3-ATAC asynchrony. The subclass identity and age of the cells are annotated beneath. **(C)** The distribution of per-cell lagging and leading states distances across various ODC maturation stages and three ages of OPCs. **(D)** The differences in remodeling rates between H3K27me3 and H3K27ac during ODC maturation are depicted here. Each line indicates the average rate differences derived from cells across the three age groups. **(E)** Heatmap showing the H3K27me3 signals of cCREs across various ODC maturation stages and three age groups of OPCs. The cCREs are organized based on their correlation with the rate of H3K27me3 remodeling. **(F)** Boxplots showing the H3K27me3 signals of cCREs across various ODC maturation stages, grouped according to their association with H3K27me3 remodeling rates. **(G)** Boxplots showing the H3K27me3 signals of cCREs across various ages of OPC and ODCs, categorized by their association with H3K27me3 remodeling rates. **(H)** Barplots showing the enrichment of high-rate association regions within each cCRE module as depicted in Figure 1G. **(I)** Genome browser view of the Prdm16 locus, a top remodeling rates-associated region showing decreased H3K27me3 signal in aged cells. **(J)** Heatmap showing the proportion of differentially modified H3K27me3 cCREs observed during aging. The figures representing significantly altered peaks are displayed on the plot. **(K)** Heatmap showing the enrichment of differentially modified H3K27me3 cCREs within DNA damage hotspots across various mouse brain cell types. **(L)** Comparison of the energy landscape between ODC maturation and in silico forced OPC transformation to astrocytes.

### Repressive epigenetic memory loss is linked to the drift of OPC fate probability

The loss of heterochromatin is considered a fundamental driver of cellular decline during aging^3,49,50^. We then analyzed changes in H3K27me3 at the modification level within 10-kb non-overlapping bins across the entire genome in ODC lineage cells. Our results showed that the most significant wave of H3K27me3 loss occurred in 43-week-old mice, while H3K27me3 gain mainly took place in 78-week-old mice (**Figure 3J**). In contrast, few regions show altered H3K27ac levels in aged cells (**Figure S3G**). We previously identified cell-type-specific hotspots of DNA damage in the mouse brain and showed that these regions are also hotspots of epigenetic erosion^51,52^. As expected, the regions exhibiting age-associated H3K27me3 alterations are enriched in OPC or ODC DNA damage hotspots when compared to the background reference mouse brain cCREs^53^ (**Figures 3K** and **S3I**). Interestingly, the cCREs in H3K27me3 erosion regions within OPCs also positively correlate with astrocyte-specific DNA damage hotspots, indicating a potential shared cellular decline phenotype between these two distinct cell types. Indeed, multiple lines of evidence support the notion that OPCs can transform into astrocytes under stress conditions^54^ or with aging^55^. To provide a thermodynamic basis for this observation, we simulated the OPC-to-astrocyte transformation by constraining the directions of cell displacement vectors and computed the H3K27me3 priming energy during the forced transition (**Figures S3J**). In young OPCs, a high activation barrier prevents progenitor cells from adopting an incorrect fate. However, in aged cells, this barrier decreases significantly, potentially shifting the balance of fate decisions toward astrocytes (**Figure 3L**). Therefore, SeqTag can provide testable hypotheses to uncover the underlying principle driving age-related functional decline.

**Figure S3.**
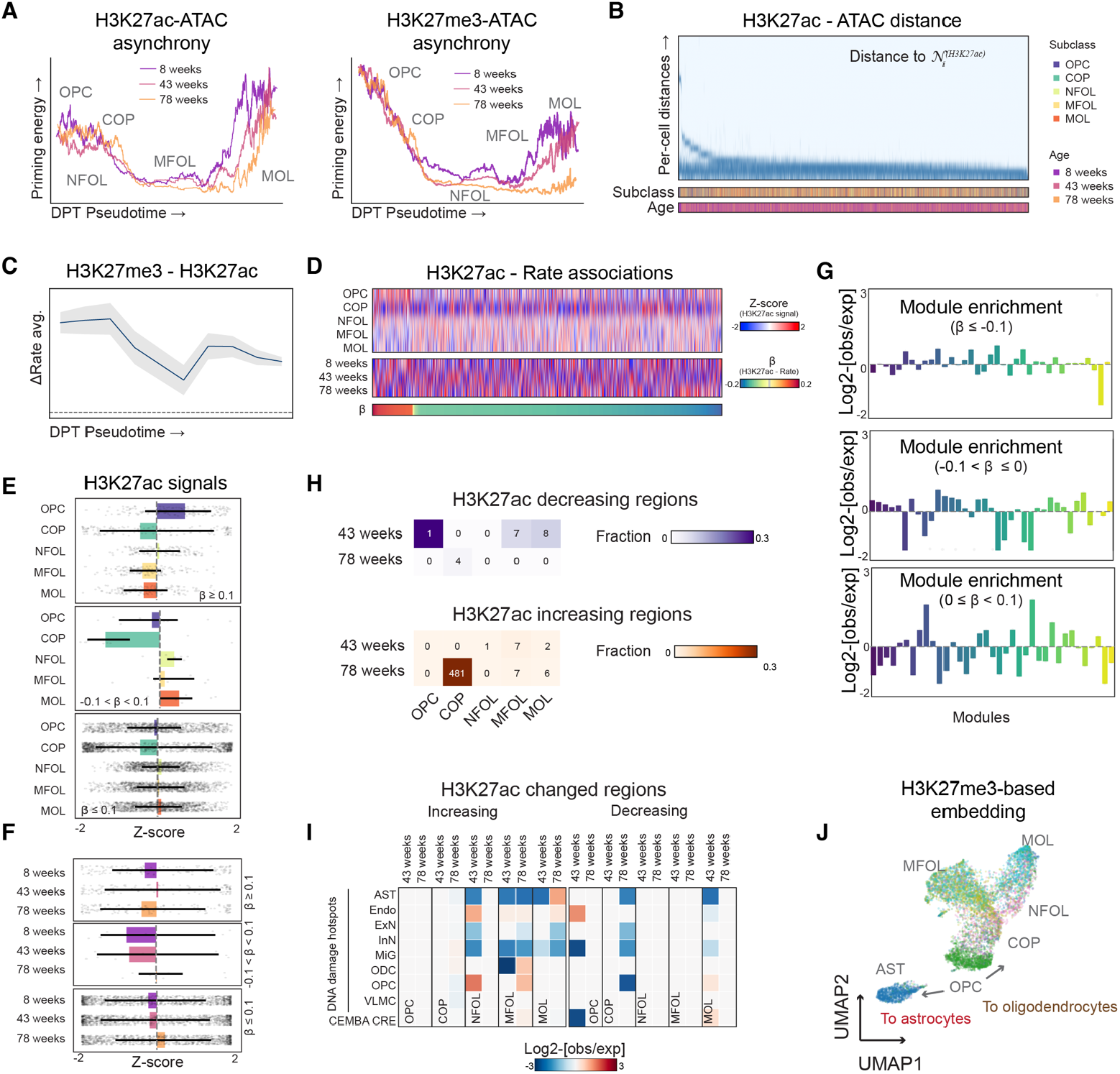
Age-associated changes in epigenetic remodeling during OPC differentiation. **(A)** Priming energy landscapes derived from H3K27ac-ATAC asynchrony (left) and H3K27me3-ATAC asynchrony (right) during ODC maturation of three ages. **(B)** Heatmap showing the normalized per-cell distance between lagging and leading states of individual cells derived from H3K27ac-ATAC asynchrony. The subclass identity and age of the cells are annotated below. **(C)** The remodeling rate disparities between H3K27me3 and H3K27ac during ODC maturation are depicted. **(D)** Heatmap showing the H3K27ac signals of cCREs across various ODC maturation stages and three age points of OPCs. The cCREs are ordered based on their correlation with the H3K27me3 remodeling rate. **(E)** Boxplots showing the H3K27ac signals of cCREs across various ODC maturation stages, categorized according to their association with H3K27ac remodeling rates. **(F)** Boxplots showing the H3K27ac signals of cCREs across various ages of OPC and ODCs, categorized by their association with H3K27ac remodeling rates. **(G)** Barplots showing the enrichment of groups regions with different rate associations across each cCRE module as depicted in Figure 1G. **(H)** Heatmap showing the proportion of differentially modified H3K27ac cCREs observed during the aging process. The plot displays the peaks with significant changes in their numbers. **(I)** Heatmap showing the enrichment of differentially modified H3K27ac cCREs within DNA damage hotspots across various mouse brain cell types. **(J)** To simulate the OPC transformation into astrocytes, astrocytes and ODC lineage cells are co-embedded to calculate the diffusion pseudotime of the two trajectories.

### The rate of cell identity loss during aging varies across different cell types

Based on the Information Theory of Aging, aging results from a gradual loss of epigenetic information, which increases regulatory entropy and leads to a decline in robust cellular identity ^3^. In line with this, we found that, for 20 of the 22 neuron subclasses, cell-type composition does not change with age (**Figure 4A**). We evaluated the regulatory entropy of single cells across different epigenetic modalities: for each cell, we assessed the dispersion of its transcriptional states when mapping its nearest epigenetic neighbors onto RNA embeddings, and the structural spread of this neighborhood reflects the thermodynamic uncertainty of the cell’s phenotype (**Figure 4B**). A broad, diffuse distribution indicates high entropy, representing active exploration of alternative cellular states under the flexible epigenetic landscape. In contrast, a tightly clustered neighborhood indicates low entropy, corresponding to lineage-restricted cells that occupy stable, low-energy states. By comparing the gradient of entropy increase among immediately related cells, we found that different cell types vary in their rate of identity decline: inhibitory neurons and vascular cells are more susceptible to this challenge, whereas astrocytes are the most resistant ones (**Figures S4A-C**). An increased proportion of high-entropy cells is a common trend across various cell types, primarily driven by chromatin accessibility and H3K27me3 drifts, with less influence from H3K27ac (**Figures 4D, 4E, S4D**, and **S4E**). We predicted the associated out-of-lineage signatures for different cell types during aging by calculating the weighted sum of the entropy-driving cCREs that overlap with cell-type-specific H3K27ac-modified regions. We discovered that the loss of H3K27me3 is linked to cell identity shifts similar to those associated with increased chromatin accessibility. For example, the loss of H3K27me3 is linked to H3K27ac regions in various non-neuron cell types, which also coincide with increased chromatin accessibility in the same cell types (**Figure 4F**). Conversely, the gain of H3K27me3 appears less associated with the reduction in chromatin accessibility (**Figure S4F**). Thus, when restrictive epigenetic barriers break down, cells lose strict transcriptional control, permitting non-lineage genes to be aberrantly expressed and leading to cell-fate drift and malfunctions.

**Figure 4.**
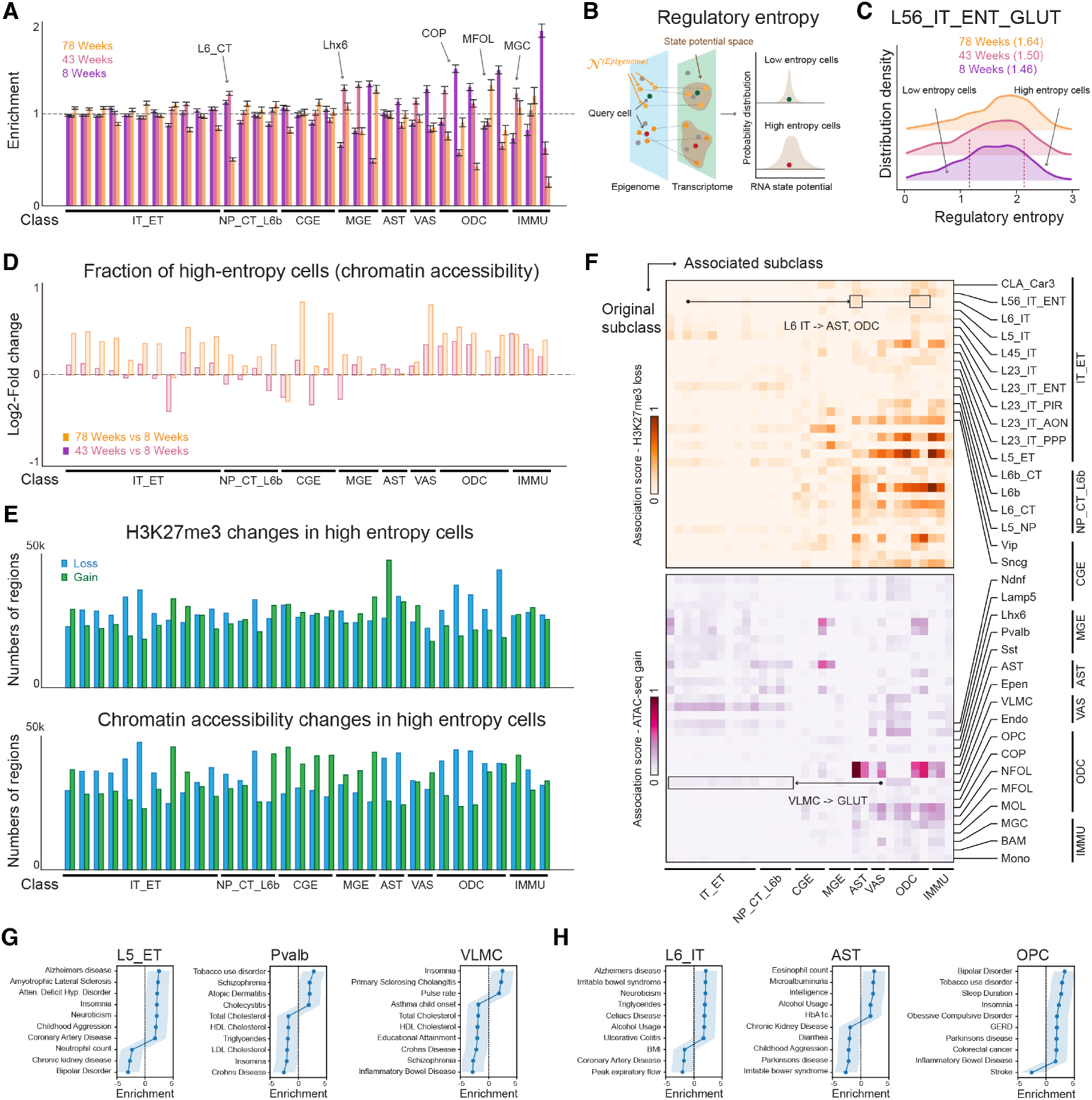
Epigenetic erosion of cell-type specific signatures in the aged brain. **(A)** Barcode plots showing the cell type proportion changes during mouse brain aging. **(B)** The regulatory entropy can be estimated by the probable distributions within the transcriptional embedding space when sharing a similar epigenetic state. The loss of cell-type-specific epigenetic signatures is proportional to the Shannon entropy of the state distribution probability. **(C)** Distribution of regulatory entropy (ATAC-seq) for L56 IT ENT GLUT neurons across different age groups. **(D)** The changes in the proportion of detected high-entropy cells (ATAC-seq) across various L2 cell subclasses throughout the aging process. **(E)** The numbers of regions exhibiting differential H3K27me3 modification (upper) and differential accessibility (bottom) in high-entropy cells across various L2 cell subclasses. **(F)** Heatmap showing the correlation between regions exhibiting H3K27me3 loss (upper panel) and accessibility gain (lower panel) in conjunction with cell-type specific H3K27ac signatures across various L2 cell subclasses. **(G)** Association of cCREs overlapped with chromatin accessibility gain regions in the human orthologs, linked to GWAS traits of diseases relevant to L5 ET GLUT, Pvalb GABA, and VLMC cell types. **(H)** Association of cCREs overlapped with H3K27me3 loss regions in human orthologs with GWAS traits related to diseases in L6 IT GLUT, AST, and OPC cell types.

**Figure S4.**
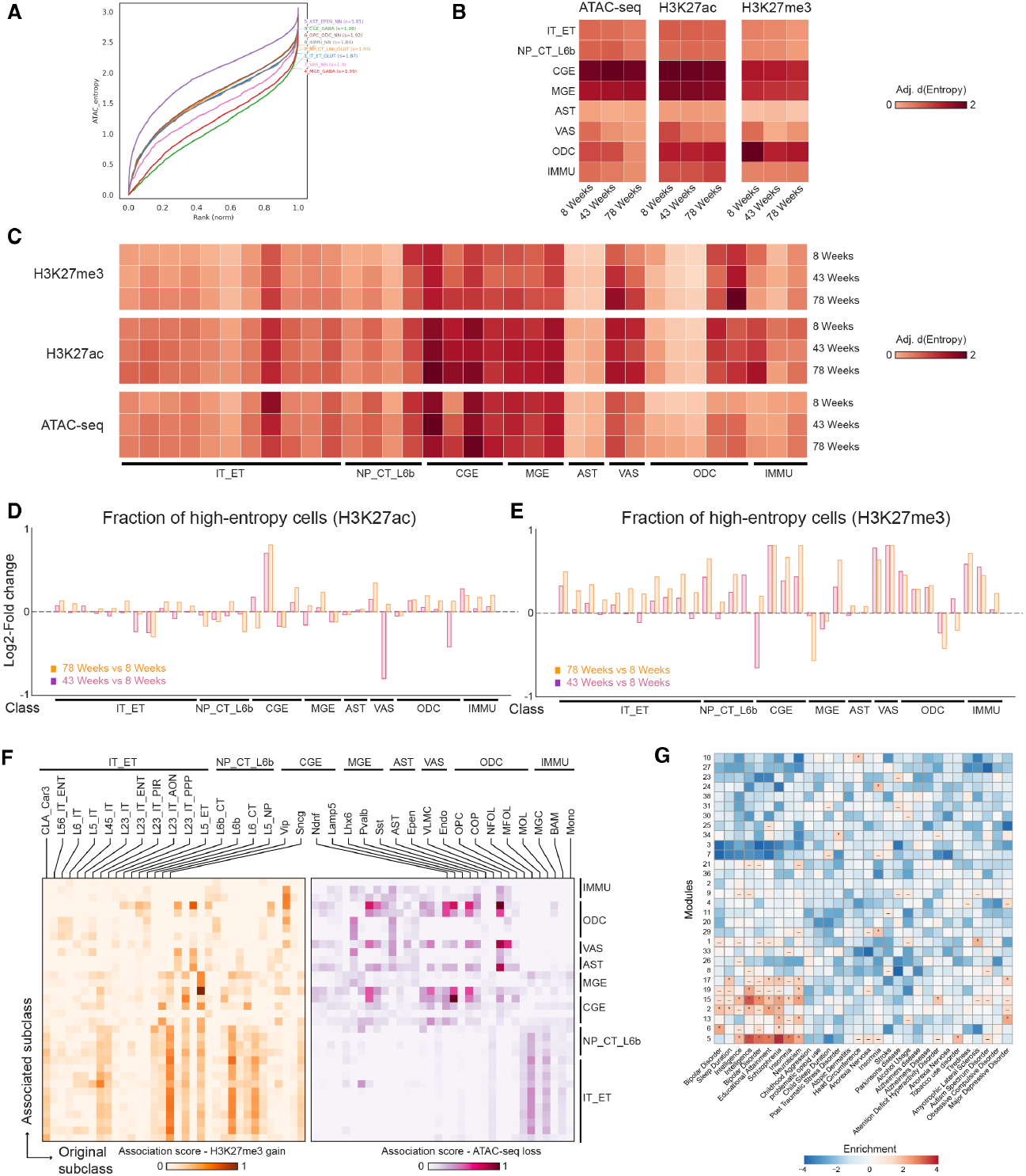
Epigenetic erosion of the aged brain cells. **(A)** Representative plots of cells ordered by ATAC-seq regulatory entropy values. Colors represent various L1 cell classes of 8-week-old mice. **(B)** Heatmap showing the estimated rates of entropy increase across various L1 cell classes, derived from ATAC-seq, H3K27ac, and H3K27me3 modalities. **(C)** Heatmap showing the estimated rates of entropy increase across various L2 cell subclasses, derived from ATAC-seq, H3K27ac, and H3K27me3 modalities. **(D)** The changes in the proportion of identified high-entropy cells (H3K27ac) across various L2 cell subclasses throughout the aging process. **(E)** The changes in the proportion of detected high-entropy cells (H3K27me3) across various L2 cell subclasses throughout the aging process. **(F)** Heatmap showing the correlation between regions of H3K27me3 gain (left) and accessibility loss (right) with cell-type-specific H3K27ac signatures across various L2 cell subclasses. **(G)** Heatmap showing the association of cCREs in the human orthologues with GWAS traits of diseases across various modules, as depicted in Figure 1G.

### Drivers of cell identity loss are associated with disease risks

Aging remains the primary risk factor for most neurodegenerative and neurological disorders, but the precise mechanistic connection between biological aging and genetic susceptibility is still not well understood. One possibility is that genetic disease risk is dynamic: in early life, non-coding risk variants are held within repressed chromatin; however, as aging causes epigenetic erosion and lowers the energy barrier, these regions may gain abnormal exposure, presenting risk loci to transcriptional machinery. To test this, we mapped the chromatin accessibility and H3K27me3 entropy-driving cCREs from our mouse model to their human orthologues and performed enrichment analyses for human GWAS traits (**Figures 4G, 4H**, and **S4G**). This analysis revealed cell-type-specific disease vulnerabilities. For example, we discovered an association between VLMC and Insomnia, in which aged VLMCs exhibit increased chromatin accessibility at enhancers of multiple excitatory neuron subclasses that are closely related to Insomnia^56^ (**Figures 4F** and **G**). Another example is that the de-repressed cCREs in L6 IT neurons are linked to Alzheimer’s Disease; these regions also show signs of two well-known AD-vulnerable cell types, astrocytes and oligodendrocytes^57^ (**Figures 4F** and **H**). Therefore, age-related epigenetic erosion weakens cell identity and raises disease risk.

## Discussion

Advances in single-cell multiomics have transformed our approach to studying gene regulation. However, existing efforts largely focus on data integration, projecting distinct molecular layers into a unified latent space to extract shared structures and catalog cell states^58^. Here, we challenge this approach: we propose the concept of molecular asynchrony, which suggests that the discordance among regulatory hierarchies is not technical noise to be regressed out; instead, it contains rich information on cellular kinetics. To measure this asynchrony, we developed SeqTag, a scalable, tri-modal single-cell technology for simultaneous analysis of chromatin accessibility, histone modification, and gene expression in single cells. With the optimized sequential tagmentation approach and instrument-free droplet-based preamplification, SeqTag minimized cross-modality signal leakages without sacrificing sensitivity. This specific tri-modal combination at single-cell resolution provides leading and lagging coordinates, which are a prerequisite for modeling the molecular asynchrony. By mapping the multimodal profiles onto a non-equilibrium thermodynamic framework, we prove that the temporal lag among regulatory layers was coded in Waddington’s landscape. This allows us to estimate cell kinetic parameters from unbiased, high-throughput single-cell data and bypasses the need for prior knowledge when analyzing complex cellular trajectories.

We applied SeqTag to the aging mouse brain to provide direct, quantitative evidence supporting the Information Theory of Aging^3^. We found that for progenitor cells (e.g., OPCs) to successfully pass the activation energy barrier, multiple regulatory layers must be primed in a coordinated manner. As cells age, the thermodynamic properties change at varying degrees for different regulatory layers. Such decoupled remodeling, primarily characterized by the loss of heterochromatin, reduces the activation barrier required to restrict cellular lineage and can create metastable intermediate states that trap aged progenitor cells during differentiation, potentially shifting the equilibrium toward alternative fates. Our thermodynamic model proposed a dynamic model of genetic susceptibility. Our data hint that in young cells, the risk loci are safely sequestered in repressed heterochromatin. However, as repressive chromatin erodes with age, such loci can gain accessibility and be exposed to the transcriptional machinery, thereby unlocking disease susceptibility.

While molecular asynchrony and SeqTag provide a robust framework for analysis of single-cell kinetics, we acknowledge the following limitations. (1) Data Sparsity and Allelic Dropout: Like other single-cell epigenomic assays, SeqTag faces inherent data sparsity, with largely binary chromatin profiles. Although our k-NN centroid projection helps reduce this issue, the priming energy calculations rely on aggregated signals across regions rather than on single-locus kinetics. (2) Thermodynamics assumption: Real-time, genome-wide tracking of epigenetic dynamics in individual cells is currently technologically infeasible. Inspired by the coupled mechanical system, we derived the kinetic parameters from a static snapshot of cell populations. However, our inference depends on the assumption of ergodic continuity within Waddington’s landscape, and catastrophic cellular events (e.g., acute ischemic stroke or viral infection) may introduce extreme momentum that falls outside the model’s predictive bounds.

In summary, we establish that the temporal lag among multiple molecular layers is essential to establishing cellular state. By modeling the molecular asynchrony, SeqTag provides a new lens for studying gene regulation, not the static changes in gene expression or chromatin states, but as dynamic shifts in the thermodynamic properties of cells.

## Methods

### Cell culture and nuclei preparation

HeLa S3 (ATCC, CCL-2.2) and NIH/3T3 (ATCC, CRL-1658) cells were maintained under standard conditions. For nuclei preparation, cells were dissociated into single-cell suspensions and harvested by centrifugation at 600 × g for 10 min at 4 °C. The resulting cell pellets were washed with PBS (ThermoFisher Scientific, 10010-23) and quantified using a Bio-Rad TC20 cell counter. To isolate the nuclei, the cells were gently resuspended to a concentration of 2,000-4,000 cells/µL in the Optimized Nuclei Isolation Lysis Buffer (OMNI Buffer)^59^. The suspension was incubated on ice for 5 mins to facilitate membrane permeabilization. Following incubation, the nuclei were recovered by centrifugation at 1,000 × g for 10 min at 4 °C. The resulting nuclei pellets were then immediately processed for subsequent SeqTag experiments.

### Mouse tissue processing and nuclei isolation

Male and female C57BL/6J mice (Strain JR#: 000664) were purchased from The Jackson Laboratory at three ages (8, 43, and 78 weeks). To prepare single-nucleus suspensions, frozen tissues were mechanically homogenized via douncing in ice-cold Complete Douncing Buffer. The resulting suspension was filtered through a 30-µm Cell-Tric filter (Sysmex) and centrifuged at 1,000 × g for 10 min at 4 °C. Following a wash with cD buffer and subsequent centrifugation, nuclei pellets were resuspended in OMNI Buffer and rotated for 5 min at 4 °C. Nuclei were counted using a cell counter before proceeding to SeqTag experiments.

### Assembly of transposomes

Double-stranded transposons were prepared by separately hybridizing three adaptor sets. The annealing reactions were conducted at 95 °C for 5 min, followed by a controlled cooling ramp to 14 °C at a rate of −0.1 °C/s. For the assembly of pA-Tn5[HM], Tn5[CA], pA-Tn5[U], and Tn5[U], 1.5 µL of each pre-annealed double-stranded transposon (50 µM) was individually complexed with 9 µL of the diluted Tn5 protein. The mixtures were incubated at room temperature for 30 mins to ensure stable transposon assembly.

### SeqTag procedures

#### Antibody staining and targeted tagmentation

Following permeabilization, nuclei were counted using a cell counter, collected by centrifugation at 700 × g for 10 min at 4 °C, and the resulting pellets were gently resuspended in 50 µL of freshly prepared Medium Salt Binding Buffer #1 (Med #1 Buffer). Targeting complexes were prepared by pre-incubating 2 µL of the primary antibody with pA-Tn5 transposomes in 18 µL of Med #1 Buffer for 2 hrs. The 22 µL of pre-assembled antibody-pA-Tn5 complexes were then added to the nuclear suspension and incubated overnight with rotation at 4 °C. Next, samples were retrieved and placed on ice. and washed with freshly prepared Medium Salt Binding Buffer #2. The transposition reaction was initiated by adding 2 µL of 250 mM MgCl^2^ (to a final concentration of 10 mM) and gently mixing. The samples were then incubated in a thermal cycler at 37 °C with shaking at 550 rpm for 30 mins. The reaction was quenched by adding 17 µL of 40 mM EDTA to chelate the Mg^2+^.

#### Open Chromatin Tagmentation

The nuclei were first resuspended in 49 µL of Tagmentation Buffer, followed by adding 0.5 µL of Tn5[CA] and 0.5 µL of Tn5[U] to the suspension. The mixture was then incubated in a thermal shaker at 37 °C and 550 rpm for 30 mins. Next, 17 µL of 40 mM EDTA was added to quench the reaction.

#### Reverse transcription (RT) reaction

Nuclei pellets were resuspended in 20 µl of reverse transcription buffer mix, and 1U/µl Maxima Reverse H Minus Reverse Transcriptase (Invitrogen, EP0751) was added. The reverse transcription was performed in a thermocycler with the following program: Step 1: 50 °C × 10 mins; Step 2: 8 °C × 12 s, 15 °C × 45 s, 20 °C × 45 s, 30 °C × 30 s, 42 °C × 2 mins, 50 °C × 5 mins, go to Step 2 for additional two times; Step 3: 50 °C × 10 mins and hold at 12 °C. After the reaction, the nuclei were transferred and pooled into 1.5-ml tubes, then immediately subjected to ligation-based combinatorial barcoding.

#### Combinatorial barcoding

Nuclei pellets were resuspended and thoroughly dispersed in 1 mL 1× NEBuffer 3.1(NEB, B6003S). The homogenized nuclei suspension was then incorporated into a master ligation mix consisting of 2,262 μL nuclease-free water, 500 μL 10× T4 DNA Ligase Buffer, 50 μL of 10 mg/mL BSA, 100 μL 10× NEBuffer 3.1, and 100 μL T4 DNA Ligase (NEB, M0202L). The resulting ligation reaction mix was aliquoted in 40 µL volumes into each well of the Barcode-plate-R02 using a multichannel pipette. The plate was then incubated at 37 °C with 300 r.p.m. agitation for 30 mins in a ThermoMixer (Eppendorf) to facilitate the first round of barcode indexing. Subsequently, 10 µL of R02-Blocking-Solution (comprising 264 µL of 100 µM Blocker-R02 oligo, 250 µL of 10× T4 Ligation Buffer, and 486 µL of ultrapure H2O) was introduced to each well to neutralize unreacted linkers. A second round of ligation was performed following the same parameters, with the addition of a Termination-Solution (264 µL of 100 µM R04 Terminator oligo, 250 µL of 0.5 M EDTA, and 236 µL of ultrapure H_2_O) to quench the reaction after the 30-min incubation. Lysis was conducted at 55 °C and 850 r.p.m. for 2 h in a ThermoMixer. Following lysis, the solution was cooled to room temperature, and the genomic material was purified using 1× SPRI beads (Beckman Coulter) and eluted in 12.5 µL of H2O. The purified DNA/cDNA libraries were stored at –20 °C or –80 °C for up to four weeks.

#### Cell lysis and cDNA/DNA purification

Following lysis, 16 µL SPRI beads were added directly to the reaction and incubated for 3 min. Following magnetization, 27 µL of supernatant was transferred to a new tube for cDNA recovery, while the bead-bound fraction was reserved for gDNA. For gDNA recovery, beads were supplemented with 12 µL XP beads, washed twice with 80% ethanol, and eluted in 14 µL nuclease-free H2O. For cDNA recovery, 15 µL XP beads were added to the previously transferred supernatant, followed by identical ethanol washes and elution in 12.5 µL H2O.

#### Amplification of DNA libraries

Index PCR was performed directly on the gDNA eluate (14 µL) by adding 2 µL TruSeq i7 indices (10 µM), 2 µL Nextera N5 indices (10 µM), and 18 µL Q5 High-Fidelity 2× Master Mix. The 36 µL reaction was amplified using the following thermal profile: 72 °C for 5 min; 98 °C for 3 min; 9–16 cycles of [98 °C for 20 s, 62 °C for 20 s, and 72 °C for 150 s]; and a final extension at 72 °C for 5 min. The resulting libraries were purified with 0.85× SPRI beads, washed twice with 80% ethanol, and eluted in 15–30 µL of nuclease-free water and stored at –20 °C.

#### Amplification of cDNA libraries

The purified cDNA (12.5 µL) was supplemented with 1.5 µL 10× TdT Buffer and 0.5 µL 1 mM dCTP, denatured at 95 °C for 5 min, and chilled on ice for 3 min. 3′ tailing was performed by adding 1 µL TdT at 37 °C for 30 min, followed by inactivation at 75 °C for 20 mins. Linear amplification was initiated by adding 15.1 µL Anchor Mix [6 µl of 5× KAPA buffer, 0.6 µl of 10mM dNTPs, 0.6 µl of 10 µM Anchor-FokI-GSH-Oligo and 0.6 µl of KAPA HiFi HS (KAPA, KK2502)] using the following profile: 98 °C for 3 mins; 16 cycles of [98 °C for 15 s, 47 °C for 60 s, 68 °C for 120 s, 47 °C for 60 s, 68 °C for 120 s]; and 72 °C for 10 mins. Subsequently, 20 µL Preamplification Mix [4µl of 5× KAPA buffer, 0.5 µl of 10 mM dNTPs, 2 µl of 10 µM PA-F and PA-R primers, 0.5 µl KAPA HiFi HS] was added directly for exponential amplification: 98 °C for 3 min; 10 cycles of [98 °C for 20 s, 65 °C for 20 s, 72 °C for 150 s]; and 72 °C for 2 min. Amplified products were purified via double-size SPRI selection (0.2×/0.65×) and eluted in 17 µL nuclease-free H2O. Following double-size SPRI selection, the purified cDNA (17 µL) was integrated into a 20 µL digestion reaction [1× CutSmart and 1 U NotI-HF] and incubated at 37 °C for 60 min. The digested product was purified using 1.25× SPRI beads and eluted in 10 µL H2O. Subsequently, the eluate was subjected to Tn5[U] tagmentation in a reaction containing 1× Tagmentation Buffer [33 mM Tris-Ac, 11 mM MgAc, 66 mM KAc, and 16% DMF] and 0.5 µL 0.05 mg/mL Tn5[U]. Tagmentation was conducted at 37 °C and 550 r.p.m. for 30 mins. The final RNA library was cleaned using the QIAquick PCR purification kit and eluted in 14 µL of 0.1× Elution Buffer, which was then supplemented directly with 2 µL TruSeq i7 indices (10 µM), 2 µL Nextera N5 indices (10 µM), and 18 µL Q5 High-Fidelity 2× Master Mix. This 36 µL reaction was amplified using the following thermal profile: 72 °C for 5 mins; 98 °C for 3 mins; 9–16 cycles of [98 °C for 20 s, 62 °C for 20 s, and 72 °C for 150 s]; and a final extension at 72 °C for 5 mins. Finally, libraries were purified with 0.85× SPRI beads, washed twice with 80% ethanol, eluted in 15–30 µL of nuclease-free water, and stored at –20 °C.

### SeqTag data analysis

#### Pre-processing of SeqTag data

Libraries were sequenced with PE 100 (Read 1) + 8 + 8 + 200 (Read 2). For Read 2, positions 1-151 were used to identify cellular barcodes and modality identifiers. The first bases of BC#1, BC#2, and BC#3 are located within the 84th–87th, 47th–50th, and 10th–13th positions of Read 2, and the exact positions were identified by matching the linker sequences. The barcode sequences were mapped to the cellular barcode reference using bowtie^60^ with the parameters: “-v 1 -m 1 –norc”. Nextera adaptor sequences were removed from 3′ of DNA and RNA libraries, and Poly-dT sequences were further removed from 3′ of RNA libraries with TrimGalore^61^. Low-quality reads with Q < 30 and length < 30 bp were excluded from downstream analysis.

#### Reads mapping

Adaptor-trimmed reads were first mapped to a mouse GRCm39 or human GRCh38 (hg38) reference genome with STAR^62^ software (v.2.6.0a) for RNA or bowtie2^63^ for DNA. Mapped DNA reads were further filtered by MAPQ > 10^64^. Duplicates were removed based on the mapped position, UMI, cellular barcode, and sub-library indices. Low-coverage nuclei (<200 transcripts and < 200 unique DNA reads) were removed from downstream analysis. Bigwig files were generated with DeepTools^65^.

#### Cell clustering

RNA bam files were converted to sparse matrix with cells as columns and genes as rows. Cell clustering based on transcriptome profiles was performed using scanpy^66^. DNA bam files were converted to fragment files with “pp.make_fragment_file” function, followed by spectral embedding with the “tl.spectral” function of snapATAC2^67^. To validate the annotation results, reference-mapping of SeqTag RNA profile to published snRNA-seq dataset^68^ was performed with MapMyCells (RRID: SCR_024672).

#### Peak calling

Peak calling was performed using MACS3^36^ with the default parameters. A cutoff of p-value < 0.05 (Benjamini-Hochberg procedure) was used to filter the results. To identify cell-type-specific peaks for all the mouse cortex cell types, we used the “tl.marker_regions” function from SnapATAC2^67^ to aggregate the DNA damage signal across cells and utilize z-scores to identify specifically enriched peaks, with a p-value cutoff = 0.1.

#### Gene-cCRE pair prediction

To link cCREs to genes, we used the SnapATAC2^67^ software. We first calculated the gene activity score from ATAC-seq modality and generated a pseudo gene-cell matrix using the “pp.make_gene_matrix” function. Next, the “tl.add_cor_scores” function was used to calculate correlations between gene activity scores and accessibility signals at cCREs. To estimate background noise levels, we shuffled the cell identities and recalculated the corresponding coaccessibility scores.

#### Motif enrichment and GO analysis

Motif enrichment for cCREs of module or bivalency groups was performed with HOMER software^69^. GO term enrichment analysis for the cCRE-linked genes was performed using the DAVID^70^ (https://davidbioinformatics.nih.gov) with default parameters.

#### GWAS enrichment

The comparison of entropy-driving cCREs with GWASs of human phenotypes was performed as in the previous study^53^. Briefly, we converted the mouse entropy-driving cCREs from GRCm39 to GRCh38 coordinates using the liftOver software with the parameter “-minMatch=0.25”^71^. Next, peaks wider than 1 kb in the human genome were excluded from further analysis. Estimation of enrichment coefficients of each trait compared to the background control was performed using the LDSC software (https://github.com/bulik/ldsc)^72^.

### Deriving the asynchrony parameters of single cells

#### Definition of community space and priming vectors

We assumed the reduced-dimension RNA expression space (***χ*** ^(*RNA*)^ ⊂ ℝ^***d***^) represents the cell’s current, realized phenotypic state. Because all cells in our data have their RNA profiles measured using the same experimental procedure, the PCA coordinates of the RNA modality were used as the reference community space, eliminating the need for cross-modal integration. The epigenomic profiles (ATAC-seq and CUT&Tag) represent “tension states” that may precede transcriptional changes. For a given cell *i*, let 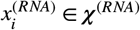 be its realized state coordinate. We identified its *k*-nearest neighbors within each modality, denoted as 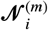. Next, we projected the epigenomic neighbors into ***χ*** ^(*RNA*)^ and calculated the local centroid (*μ*_*m*_) and the displacement vector (𝒱^(*m*)^) for each modality :

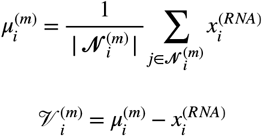

The gross magnitude of epigenetic drift is:

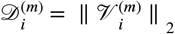

Since the RNA space contains both inherent biological and technical variance, minor displacements are not biologically meaningful. We define a baseline dispersion threshold (*α*) based on the standard deviation of the Euclidean distances to the cell’s closest RNA neighbors:

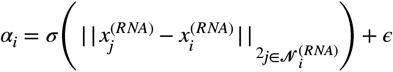

To understand how the two epigenomic layers interact, we isolated the movement of the CUT&Tag modality that aligns with the ATAC-seq modality by calculating the cosine similarity between the two epigenomic shifts:

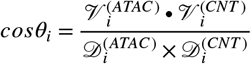

The effective unrealized space for state change beyond the baseline noise is then obtained:

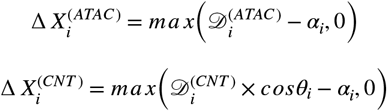

#### Epigenetic tension potential (Priming Energy, Δ *E*_*pri*_)

We modeled the cell state transition lag between layers as a coupled mechanical system. The ‘unrealized potential’ gap is the priming distance by which ATAC-seq has outpaced CUT&Tag:

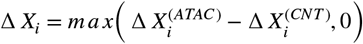

According to Hooke’s Law, the potential energy of the lagging system is: 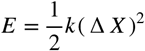. However, if the lagging modality has not moved at all (Δ *X*_2_ = 0), the coupling stiffness (*k*) is effectively zero, and no energy can be transferred. We assume the coupling stiffness scales with the ratio of their effective priming distance:

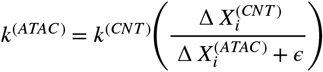

We set a global scaling parameter 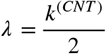 and derived the Priming Energy:

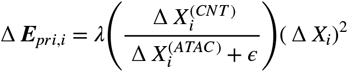

#### Cell state plasticity (Regulatory Entropy, 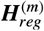)

We assumed that high variance in the projected states of a cell’s epigenomic neighbors indicates a broad, permissive epigenetic landscape, which translates to high cell-state plasticity. For density estimation, we first mean-center the neighbor coordinates and project them onto their first principal component (𝒵^(*m*)^) using Singular Value Decomposition (SVD). Next, we estimated the continuous probability density function (*P*^(*m*)^) of these projected neighbors using a Gaussian Kernel Density Estimate (KDE) with a bandwidth of 10^−3^. Finally, we calculated the Shannon entropy of the distribution to represent cell state plasticity of each epigenetic modality :

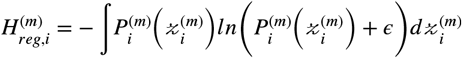

#### Epigenetic remodeling rate (*v*_*diff*_)

To detect asynchronous jumps, we define a cell’s immediate future state as its projected position on the leading ATAC-seq manifold. For a given cell, we projected its nearest neighbors strictly within the CUT&Tag space 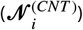 into the ATAC-seq spectral embedding coordinates (***χ*** ^(*ATAC*)^). Next, we calculated the Euclidean distance from cell to each of the 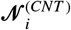 within the ***χ*** ^(*ATAC*)^:

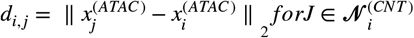

and then isolated the relative distribution of neighbors by applying a per-cell Min-Max normalization:

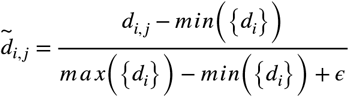

To determine if a cell is actively transitioning between states, we test the normalized distance array for bimodality. For a transitioning (bimodal) cell, the two peaks represent the “lagging” and “leading” states; the gap between the two states represents the distance (Δ*x*) that the CUT&Tag modality must traverse to synchronize with the ATAC-seq modality over the next unit of time (Δ*t*):

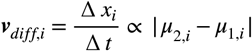

## Data availability

Raw sequencing and processed data generated in this study are available from GEO: GSE333552.

## Code availability

The customized codes used in this study are available on GitHub (https://github.com/czhulab/asynchrony).

## Acknowledgements

We thank QB3 MacroLab for the protein A-Tn5 and Tn5 enzymes. C.Z. is supported by Weill Cornell Medicine and New York Genome Center startup funds, National Institutes of Health (NIH)/National Institute of General Medical Sciences (grant no. DP2GM154011), NIH/National Cancer Institute (grant no. R61CA309705), NIH/National Human Genome Research Institute (grant no. RM1HG011014), and the MacMillan Center for the Study of the Noncoding Cancer Genome at the New York Genome Center.

## Author contributions

B.F. and C.Z. conceived the study. B.F. developed the SeqTag protocol and generated the single-cell data with the help of R.T., X.B., and D.B. C.Z. conceptualized the molecular asynchrony framework, incorporating J.S.’s input, and performed data analysis with the help of B.F. and Z.C. B.F., J.S., and C.Z. wrote the manuscript and discussed it with all authors. C.Z. supervised the study.

## Ethics declarations

The authors declared no conflict of interest.

## References

1. Waddington, C.H. (1957). The Strategy of the Genes: A Discussion of Some Aspects of Theoretical Biology (George Allen & Unwin).

2. Moris, N., Pina, C., and Arias, A.M. (2016). Transition states and cell fate decisions in epigenetic landscapes. Nat Rev Genet 17, 693–703. 10.1038/nrg.2016.98.

3. Lu, Y.R., Tian, X., and Sinclair, D.A. (2023). The Information Theory of Aging. Nat Aging 3, 1486–1499. 10.1038/s43587-023-00527-6.

4. Longo, S.K., Guo, M.G., Ji, A.L., and Khavari, P.A. (2021). Integrating single-cell and spatial transcriptomics to elucidate intercellular tissue dynamics. Nat Rev Genet 22, 627–644. 10.1038/s41576-021-00370-8.

5. Haghverdi, L., Buttner, M., Wolf, F.A., Buettner, F., and Theis, F.J. (2016). Diffusion pseudotime robustly reconstructs lineage branching. Nat Methods 13, 845–848. 10.1038/nmeth.3971.

6. Street, K., Risso, D., Fletcher, R.B., Das, D., Ngai, J., Yosef, N., Purdom, E., and Dudoit, S. (2018). Slingshot: cell lineage and pseudotime inference for single-cell transcriptomics. BMC Genomics 19, 477. 10.1186/s12864-018-4772-0.

7. Trapnell, C., Cacchiarelli, D., Grimsby, J., Pokharel, P., Li, S., Morse, M., Lennon, N.J., Livak, K.J., Mikkelsen, T.S., and Rinn, J.L. (2014). The dynamics and regulators of cell fate decisions are revealed by pseudotemporal ordering of single cells. Nat Biotechnol 32, 381–386. 10.1038/nbt.2859.

8. Bergen, V., Lange, M., Peidli, S., Wolf, F.A., and Theis, F.J. (2020). Generalizing RNA velocity to transient cell states through dynamical modeling. Nat Biotechnol 38, 1408–1414. 10.1038/s41587-020-0591-3.

9. Bergen, V., Soldatov, R.A., Kharchenko, P.V., and Theis, F.J. (2021). RNA velocity-current challenges and future perspectives. Mol Syst Biol 17, e10282. 10.15252/msb.202110282.

10. La Manno, G., Soldatov, R., Zeisel, A., Braun, E., Hochgerner, H., Petukhov, V., Lidschreiber, K., Kastriti, M.E., Lonnerberg, P., Furlan, A., et al. (2018). RNA velocity of single cells. Nature 560, 494–498. 10.1038/s41586-018-0414-6.

11. Schiebinger, G., Shu, J., Tabaka, M., Cleary, B., Subramanian, V., Solomon, A., Gould, J., Liu, S., Lin, S., Berube, P., et al. (2019). Optimal-Transport Analysis of Single-Cell Gene Expression Identifies Developmental Trajectories in Reprogramming. Cell 176, 928–943 e922. 10.1016/j.cell.2019.01.006.

12. Aibar, S., Gonzalez-Blas, C.B., Moerman, T., Huynh-Thu, V.A., Imrichova, H., Hulselmans, G., Rambow, F., Marine, J.C., Geurts, P., Aerts, J., et al. (2017). SCENIC: single-cell regulatory network inference and clustering. Nat Methods 14, 1083–1086. 10.1038/nmeth.4463.

13. Ma, S., Zhang, B., LaFave, L.M., Earl, A.S., Chiang, Z., Hu, Y., Ding, J., Brack, A., Kartha, V.K., Tay, T., et al. (2020). Chromatin Potential Identified by Shared Single-Cell Profiling of RNA and Chromatin. Cell 183, 1103–1116 e1120. 10.1016/j.cell.2020.09.056.

14. Bai, D., Zhang, X., Xiang, H., Guo, Z., Zhu, C., and Yi, C. (2025). Simultaneous single-cell analysis of 5mC and 5hmC with SIMPLE-seq. Nat Biotechnol 43, 85–96. 10.1038/s41587-024-02148-9.

15. Eyring, H. (1935). The Activated Complex and the Absolute Rate of Chemical Reactions. Chemical Reviews 17, 65–77. 10.1021/cr60056a006.

16. Genuth, M., Kojima, Y., Jülich, D., Kiryu, H., and Holley, S.A. (2021). Ergodic patterns of cell state transitions underlie the reproducibility of embryonic development. bioRxiv, 2021.2008.2023.457360. 10.1101/2021.08.23.457360.

17. Weinreb, C., Wolock, S., Tusi, B.K., Socolovsky, M., and Klein, A.M. (2018). Fundamental limits on dynamic inference from single-cell snapshots. Proc Natl Acad Sci U S A 115, E2467–E2476. 10.1073/pnas.1714723115.

18. Spivakov, M., and Fisher, A.G. (2007). Epigenetic signatures of stem-cell identity. Nat Rev Genet 8, 263–271. 10.1038/nrg2046.

19. Koche, R.P., Smith, Z.D., Adli, M., Gu, H., Ku, M., Gnirke, A., Bernstein, B.E., and Meissner, A. (2011). Reprogramming factor expression initiates widespread targeted chromatin remodeling. Cell Stem Cell 8, 96–105. 10.1016/j.stem.2010.12.001.

20. Stergachis, A.B., Neph, S., Reynolds, A., Humbert, R., Miller, B., Paige, S.L., Vernot, B., Cheng, J.B., Thurman, R.E., Sandstrom, R., et al. (2013). Developmental fate and cellular maturity encoded in human regulatory DNA landscapes. Cell 154, 888–903. 10.1016/j.cell.2013.07.020.

21. Gibbs, J.W. (1902). Elementary Principles in Statistical Mechanics: Developed with Especial Reference to the Rational Foundation of Thermodynamics (Charles Scribner’s Sons).

22. Boltzmann, L. (1871). Einige allgemeine Sätze über das Wärmegleichgewicht. Sitzungsberichte der Kaiserlichen Akademie der Wissenschaften in Wien, Mathematisch-Naturwissenschaftliche Classe 63, 679–711.

23. Huang, S., Guo, Y.P., May, G., and Enver, T. (2007). Bifurcation dynamics in lineage-commitment in bipotent progenitor cells. Dev Biol 305, 695–713. 10.1016/j.ydbio.2007.02.036.

24. Buenrostro, J.D., Giresi, P.G., Zaba, L.C., Chang, H.Y., and Greenleaf, W.J. (2013). Transposition of native chromatin for fast and sensitive epigenomic profiling of open chromatin, DNA-binding proteins and nucleosome position. Nat Methods 10, 1213–1218. 10.1038/nmeth.2688.

25. Rosenberg, A.B., Roco, C.M., Muscat, R.A., Kuchina, A., Sample, P., Yao, Z., Graybuck, L.T., Peeler, D.J., Mukherjee, S., Chen, W., et al. (2018). Single-cell profiling of the developing mouse brain and spinal cord with split-pool barcoding. Science 360, 176–182. 10.1126/science.aam8999.

26. Kaya-Okur, H.S., Wu, S.J., Codomo, C.A., Pledger, E.S., Bryson, T.D., Henikoff, J.G., Ahmad, K., and Henikoff, S. (2019). CUT&Tag for efficient epigenomic profiling of small samples and single cells. Nat Commun 10, 1930. 10.1038/s41467-019-09982-5.

27. Meers, M.P., Llagas, G., Janssens, D.H., Codomo, C.A., and Henikoff, S. (2023). Multifactorial profiling of epigenetic landscapes at single-cell resolution using MulTI-Tag. Nat Biotechnol 41, 708–716. 10.1038/s41587-022-01522-9.

28. Mereu, E., Lafzi, A., Moutinho, C., Ziegenhain, C., McCarthy, D.J., Alvarez-Varela, A., Batlle, E., Sagar Grun, D., Lau, J.K., et al. (2020). Benchmarking single-cell RNA-sequencing protocols for cell atlas projects. Nat Biotechnol 38, 747–755. 10.1038/s41587-020-0469-4.

29. Clark, I.C., Fontanez, K.M., Meltzer, R.H., Xue, Y., Hayford, C., May-Zhang, A., D’Amato, C., Osman, A., Zhang, J.Q., Hettige, P., et al. (2023). Microfluidics-free single-cell genomics with templated emulsification. Nat Biotechnol 41, 1557–1566. 10.1038/s41587-023-01685-z.

30. Cao, J., Cusanovich, D.A., Ramani, V., Aghamirzaie, D., Pliner, H.A., Hill, A.J., Daza, R.M., McFaline-Figueroa, J.L., Packer, J.S., Christiansen, L., et al. (2018). Joint profiling of chromatin accessibility and gene expression in thousands of single cells. Science 361, 1380–1385. 10.1126/science.aau0730.

31. Zhu, C., Yu, M., Huang, H., Juric, I., Abnousi, A., Hu, R., Lucero, J., Behrens, M.M., Hu, M., and Ren, B. (2019). An ultra high-throughput method for single-cell joint analysis of open chromatin and transcriptome. Nat Struct Mol Biol 26, 1063–1070. 10.1038/s41594-019-0323-x.

32. Chen, S., Lake, B.B., and Zhang, K. (2019). High-throughput sequencing of the transcriptome and chromatin accessibility in the same cell. Nat Biotechnol 37, 1452–1457. 10.1038/s41587-019-0290-0.

33. Li, G., Liu, Y., Zhang, Y., Kubo, N., Yu, M., Fang, R., Kellis, M., and Ren, B. (2019). Joint profiling of DNA methylation and chromatin architecture in single cells. Nat Methods 16, 991–993. 10.1038/s41592-019-0502-z.

34. Xu, W., Yang, W., Zhang, Y., Chen, Y., Hong, N., Zhang, Q., Wang, X., Hu, Y., Song, K., Jin, W., and Chen, X. (2022). ISSAAC-seq enables sensitive and flexible multimodal profiling of chromatin accessibility and gene expression in single cells. Nat Methods 19, 1243–1249. 10.1038/s41592-022-01601-4.

35. Yao, Z., van Velthoven, C.T.J., Kunst, M., Zhang, M., McMillen, D., Lee, C., Jung, W., Goldy, J., Abdelhak, A., Aitken, M., et al. (2023). A high-resolution transcriptomic and spatial atlas of cell types in the whole mouse brain. Nature 624, 317–332. 10.1038/s41586-023-06812-z.

36. Zhang, Y., Liu, T., Meyer, C.A., Eeckhoute, J., Johnson, D.S., Bernstein, B.E., Nusbaum, C., Myers, R.M., Brown, M., Li, W., and Liu, X.S. (2008). Model-based analysis of ChIP-Seq (MACS). Genome Biol 9, R137. 10.1186/gb-2008-9-9-r137.

37. Pozniak, C.D., Langseth, A.J., Dijkgraaf, G.J., Choe, Y., Werb, Z., and Pleasure, S.J. (2010). Sox10 directs neural stem cells toward the oligodendrocyte lineage by decreasing Suppressor of Fused expression. Proc Natl Acad Sci U S A 107, 21795–21800. 10.1073/pnas.1016485107.

38. Anderson, S.A., Eisenstat, D.D., Shi, L., and Rubenstein, J.L. (1997). Interneuron migration from basal forebrain to neocortex: dependence on Dlx genes. Science 278, 474–476. 10.1126/science.278.5337.474.

39. Guo, S., Wang, H., and Yin, Y. (2022). Microglia Polarization From M1 to M2 in Neurodegenerative Diseases. Front Aging Neurosci 14, 815347. 10.3389/fnagi.2022.815347.

40. Pekny, M., and Nilsson, M. (2005). Astrocyte activation and reactive gliosis. Glia 50, 427–434. 10.1002/glia.20207.

41. Wang, Y., Xiao, M., Chen, X., Chen, L., Xu, Y., Lv, L., Wang, P., Yang, H., Ma, S., Lin, H., et al. (2015). WT1 recruits TET2 to regulate its target gene expression and suppress leukemia cell proliferation. Mol Cell 57, 662–673. 10.1016/j.molcel.2014.12.023.

42. Pols, M.S., van Meel, E., Oorschot, V., ten Brink, C., Fukuda, M., Swetha, M.G., Mayor, S., and Klumperman, J. (2013). hVps41 and VAMP7 function in direct TGN to late endosome transport of lysosomal membrane proteins. Nat Commun 4, 1361. 10.1038/ncomms2360.

43. Sahara, S., Aoto, M., Eguchi, Y., Imamoto, N., Yoneda, Y., and Tsujimoto, Y. (1999). Acinus is a caspase-3-activated protein required for apoptotic chromatin condensation. Nature 401, 168–173. 10.1038/43678.

44. Ziskin, J.L., Nishiyama, A., Rubio, M., Fukaya, M., and Bergles, D.E. (2007). Vesicular release of glutamate from unmyelinated axons in white matter. Nat Neurosci 10, 321–330. 10.1038/nn1854.

45. Guerrier, S., Coutinho-Budd, J., Sassa, T., Gresset, A., Jordan, N.V., Chen, K., Jin, W.L., Frost, A., and Polleux, F. (2009). The F-BAR domain of srGAP2 induces membrane protrusions required for neuronal migration and morphogenesis. Cell 138, 990–1004. 10.1016/j.cell.2009.06.047.

46. Fujita, Y., Shirataki, H., Sakisaka, T., Asakura, T., Ohya, T., Kotani, H., Yokoyama, S., Nishioka, H., Matsuura, Y., Mizoguchi, A., et al. (1998). Tomosyn: a syntaxin-1-binding protein that forms a novel complex in the neurotransmitter release process. Neuron 20, 905–915. 10.1016/s0896-6273(00)80472-9.

47. Chuikov, S., Levi, B.P., Smith, M.L., and Morrison, S.J. (2010). Prdm16 promotes stem cell maintenance in multiple tissues, partly by regulating oxidative stress. Nat Cell Biol 12, 999–1006. 10.1038/ncb2101.

48. Zong, H., Parada, L.F., and Baker, S.J. (2015). Cell of origin for malignant gliomas and its implication in therapeutic development. Cold Spring Harb Perspect Biol 7. 10.1101/cshperspect.a020610.

49. Villeponteau, B. (1997). The heterochromatin loss model of aging. Exp Gerontol 32, 383–394. 10.1016/s0531-5565(96)00155-6.

50. Yang, J.H., Hayano, M., Griffin, P.T., Amorim, J.A., Bonkowski, M.S., Apostolides, J.K., Salfati, E.L., Blanchette, M., Munding, E.M., Bhakta, M., et al. (2023). Loss of epigenetic information as a cause of mammalian aging. Cell 186, 305–326 e327. 10.1016/j.cell.2022.12.027.

51. Bai, D., Cao, Z., Attada, N., Song, J., and Zhu, C. (2025). Single-cell parallel analysis of DNA damage and transcriptome reveals selective genome vulnerability. Nat Methods 22, 962–972. 10.1038/s41592-025-02632-3.

52. Zhang, Y., Amaral, M.L., Zhu, C., Grieco, S.F., Hou, X., Lin, L., Buchanan, J., Tong, L., Preissl, S., Xu, X., and Ren, B. (2022). Single-cell epigenome analysis reveals age-associated decay of heterochromatin domains in excitatory neurons in the mouse brain. Cell Res 32, 1008–1021. 10.1038/s41422-022-00719-6.

53. Li, Y.E., Preissl, S., Hou, X., Zhang, Z., Zhang, K., Qiu, Y., Poirion, O.B., Li, B., Chiou, J., Liu, H., et al. (2021). An atlas of gene regulatory elements in adult mouse cerebrum. Nature 598, 129–136. 10.1038/s41586-021-03604-1.

54. Bai, X., Zhao, N., Koupourtidou, C., Fang, L.P., Schwarz, V., Caudal, L.C., Zhao, R., Hirrlinger, J., Walz, W., Bian, S., et al. (2023). In the mouse cortex, oligodendrocytes regain a plastic capacity, transforming into astrocytes after acute injury. Dev Cell 58, 1153–1169 e1155. 10.1016/j.devcel.2023.04.016.

55. Baror, R., Neumann, B., Segel, M., Chalut, K.J., Fancy, S.P.J., Schafer, D.P., and Franklin, R.J.M. (2019). Transforming growth factor-beta renders ageing microglia inhibitory to oligodendrocyte generation by CNS progenitors. Glia 67, 1374–1384. 10.1002/glia.23612.

56. Stein, M.B., McCarthy, M.J., Chen, C.Y., Jain, S., Gelernter, J., He, F., Heeringa, S.G., Kessler, R.C., Nock, M.K., Ripke, S., et al. (2018). Genome-wide analysis of insomnia disorder. Mol Psychiatry 23, 2238–2250. 10.1038/s41380-018-0033-5.

57. Mathys, H., Davila-Velderrain, J., Peng, Z., Gao, F., Mohammadi, S., Young, J.Z., Menon, M., He, L., Abdurrob, F., Jiang, X., et al. (2019). Single-cell transcriptomic analysis of Alzheimer’s disease. Nature 570, 332–337. 10.1038/s41586-019-1195-2.

58. Argelaguet, R., Cuomo, A.S.E., Stegle, O., and Marioni, J.C. (2021). Computational principles and challenges in single-cell data integration. Nat Biotechnol 39, 1202–1215. 10.1038/s41587-021-00895-7.

59. Corces, M.R., Trevino, A.E., Hamilton, E.G., Greenside, P.G., Sinnott-Armstrong, N.A., Vesuna, S., Satpathy, A.T., Rubin, A.J., Montine, K.S., Wu, B., et al. (2017). An improved ATAC-seq protocol reduces background and enables interrogation of frozen tissues. Nat Methods 14, 959–962. 10.1038/nmeth.4396.

60. Langmead, B., Trapnell, C., Pop, M., and Salzberg, S.L. (2009). Ultrafast and memory-efficient alignment of short DNA sequences to the human genome. Genome Biol 10, R25. 10.1186/gb-2009-10-3-r25.

61. Krueger, F. (2015). Trim galore. A wrapper tool around Cutadapt and FastQC to consistently apply quality and adapter trimming to FastQ files 516, 517.

62. Dobin, A., Davis, C.A., Schlesinger, F., Drenkow, J., Zaleski, C., Jha, S., Batut, P., Chaisson, M., and Gingeras, T.R. (2013). STAR: ultrafast universal RNA-seq aligner. Bioinformatics 29, 15–21. 10.1093/bioinformatics/bts635.

63. Langmead, B., and Salzberg, S.L. (2012). Fast gapped-read alignment with Bowtie 2. Nat Methods 9, 357–359. 10.1038/nmeth.1923.

64. Li, H., Handsaker, B., Wysoker, A., Fennell, T., Ruan, J., Homer, N., Marth, G., Abecasis, G., Durbin, R., and Genome Project Data Processing, S. (2009). The Sequence Alignment/Map format and SAMtools. Bioinformatics 25, 2078–2079. 10.1093/bioinformatics/btp352.

65. Ramirez, F., Ryan, D.P., Gruning, B., Bhardwaj, V., Kilpert, F., Richter, A.S., Heyne, S., Dundar, F., and Manke, T. (2016). deepTools2: a next generation web server for deep-sequencing data analysis. Nucleic Acids Res 44, W160–165. 10.1093/nar/gkw257.

66. Wolf, F.A., Angerer, P., and Theis, F.J. (2018). SCANPY: large-scale single-cell gene expression data analysis. Genome Biol 19, 15. 10.1186/s13059-017-1382-0.

67. Zhang, K., Zemke, N.R., Armand, E.J., and Ren, B. (2024). A fast, scalable and versatile tool for analysis of single-cell omics data. Nat Methods 21, 217–227. 10.1038/s41592-023-02139-9.

68. Yao, Z., Liu, H., Xie, F., Fischer, S., Adkins, R.S., Aldridge, A.I., Ament, S.A., Bartlett, A., Behrens, M.M., Van den Berge, K., et al. (2021). A transcriptomic and epigenomic cell atlas of the mouse primary motor cortex. Nature 598, 103–110. 10.1038/s41586-021-03500-8.

69. Heinz, S., Benner, C., Spann, N., Bertolino, E., Lin, Y.C., Laslo, P., Cheng, J.X., Murre, C., Singh, H., and Glass, C.K. (2010). Simple combinations of lineage-determining transcription factors prime cis-regulatory elements required for macrophage and B cell identities. Mol Cell 38, 576–589. 10.1016/j.molcel.2010.05.004.

70. Huang da, W., Sherman, B.T., and Lempicki, R.A. (2009). Systematic and integrative analysis of large gene lists using DAVID bioinformatics resources. Nat Protoc 4, 44–57. 10.1038/nprot.2008.211.

71. Lee, B.T., Barber, G.P., Benet-Pages, A., Casper, J., Clawson, H., Diekhans, M., Fischer, C., Gonzalez, J.N., Hinrichs, A.S., Lee, C.M., et al. (2022). The UCSC Genome Browser database: 2022 update. Nucleic Acids Res 50, D1115–D1122. 10.1093/nar/gkab959.

72. Bulik-Sullivan, B.K., Loh, P.R., Finucane, H.K., Ripke, S., Yang, J., Schizophrenia Working Group of the Psychiatric Genomics, C., Patterson, N., Daly, M.J., Price, A.L., and Neale, B.M. (2015). LD Score regression distinguishes confounding from polygenicity in genome-wide association studies. Nat Genet 47, 291–295. 10.1038/ng.3211.

